# Exosome-Derived Proteomic Signatures Highlight Pathogenic Mechanisms in Moyamoya Disease

**DOI:** 10.64898/2026.07.30.741716

**Authors:** Tulika Gupta, Ranjana Bharti, Veena Devi, Munish Kumar, Ashish Aggarwal, Jaswinder Singh Maras

## Abstract

**Background:** The etiology and molecular mechanisms of Moyamoya disease (MMD) remain unclear. Exosomes, as carriers of bioactive molecules, may reflect disease-specific alterations and serve as potential biomarkers. This study aimed to investigate disease mechanisms using proteomic profiling of serum-derived exosomes (SDEs) in MMD.

**Materials and Methods:** Peripheral blood each from 15 MMD patients and 15 healthy-controls were used to isolate SDEs via ultracentrifugation. Proteins from pooled SDEs were extracted, digested, and analyzed by LC-MS/MS. Differentially expressed proteins were examined using MetaboAnalyst, DAVID, Enrichr, STRING, and Cytoscape. Key targets were validated at transcript and protein levels using RT-qPCR and ELISA in independent cohorts.

**Results:** A total of 2,554 proteins were identified, with 213 showing differential expression (118 upregulated, 95 downregulated; p ≤ 0.05). Functional and pathway analyses revealed enrichment in angiogenesis, cytoskeletal remodeling, and endothelial signaling. PRKG2 and MYC were upregulated, while RHOA was downregulated, highlighting their involvement in focal adhesion and PI3K-AKT pathways. Validation confirmed these findings.

**Conclusion:** Dysregulated proteins were linked to RHOA-ROCK and PI3K-Akt signaling, suggesting their role in driving VSMC phenotypic switching, contributing to vascular occlusion. These findings indicate that altered exosomal-proteins may participate in maladaptive vascular remodeling, although the initial trigger for VSMC transition remains unknown.

## Introduction

Moyamoya disease is a rare and idiopathic cerebrovascular disorder, characterized by progressive, non-atherosclerotic, often bilateral narrowing of the terminal portion of the internal carotid arteries (ICAs) and the circle of Willis [1]. This narrowing leads to the development of abnormal, compensatory collateral vessels, referred to as moyamoya vessels, which have a hazy, smoke like appearance on radiographs [2]. Clinically, Moyamoya disease presents with ischemic events such as transient ischemic attacks (TIAs) and ischemic strokes predominantly in children, whereas adults are more frequently affected by intracranial haemorrhages [3]. At present, revascularization surgery is the primary treatment for MMD, but there are no effective therapies targeting the pathogenesis [4].

Cerebrovascular occlusion in Moyamoya disease is primarily attributed to progressive fibro-cellular intimal thickening and structural disruption of the internal elastic lamina, which together lead to significant narrowing or complete blockage of intracranial arteries and impaired cerebral perfusion [5, 6]. Histopathological analysis of affected vessels, particularly those within the circle of Willis and its major branches, reveals hallmark features including dense intimal hyperplasia, duplication and fragmentation of the internal elastic lamina, and its characteristic wavy or disrupted morphology. These changes are accompanied by a reduction in the external arterial diameter and pronounced thinning of the medial layer, reflecting a global remodeling of the vessel wall architecture [6]. Immunohistochemical analysis have shown that the thickened intima predominantly comprises vascular smooth muscle cells (VSMCs) that have undergone phenotypic switching from a contractile to a synthetic state [7, 8]. The presence of proliferating cell nuclear antigen (PCNA)-positive VSMCs within the intima further suggests that active proliferation and phenotypic modulation are key drivers of neointimal formation [9]. Complementing these changes, apoptosis-mediated thinning of the medial layer further undermines vessel wall integrity, creating a permissive environment for stenosis progression [7, 8]. Together, these pathological alterations are thought to contribute to the progressive arterial narrowing characteristic of Moyamoya disease [10]. This dynamic process is tightly regulated by complex signalling networks, many of which are modulated by exosome-mediated intercellular communication [11].

Recently, exosomes have garnered significant attention for their potential as both diagnostic biomarkers and therapeutic agents. Exosomes are nanosized extracellular vesicles secreted by nearly all cell types and present in diverse body fluids. They carry bioactive cargo, including proteins, lipids, and nucleic acids, and act as mediators of intercellular communication, thereby reflecting the physiological or pathological state of their cells of origin [12]. As key mediators of intercellular communication, they influence recipient cell behaviour and are crucial in maintaining tissue homeostasis and mediating disease progression [13]. In neurodegenerative diseases, for example, exosomes derived from activated glial cells have been shown to transfer neurotoxic proteins such as amyloid-β and α-synuclein, contributing to the pathogenesis of Alzheimer’s and Parkinson’s diseases, respectively [14]. Within the tumour microenvironment, exosomes facilitate cancer progression by promoting angiogenesis, immune escape, and metastasis [15]. Importantly, in the vascular system, exosomes regulate key processes such as angiogenesis, inflammation, and extracellular matrix (ECM) remodeling, thereby highlighting their potential relevance in cerebrovascular disorders like Moyamoya disease [16, 17].

Given their role as molecular surrogates of cellular states, we hypothesized that alterations in exosomal protein profiles could mirror the underlying pathophysiological mechanisms of Moyamoya disease. Accordingly, this study sought to systematically characterize serum-derived exosomal proteins from patients with MMD, with the goal of identifying their potential contributions to disease pathogenesis.

## Materials and methods

### Study design

This study was conducted in two sequential phases: discovery and validation. In the discovery phase, proteomic analysis of serum-derived exosomes from Moyamoya disease (MMD) patients and matched controls was performed using LC-MS/MS on a Q-Exactive™ Plus instrument (Thermo Fisher Scientific) to identify proteins potentially contributing to disease pathogenesis. In the validation phase, an independent cohort of MMD patients and controls was examined to validate the findings from the discovery phase. Selected candidate proteins were assessed at both the transcript level, using RT-qPCR, and the protein level, using ELISA on serum-derived exosomes, to establish their biological relevance.

### Patient selection, sample collection and storage

This study was conducted in the Department of Anatomy, Postgraduate Institute of Medical Education and Research (PGIMER), Chandigarh, India, with ethical approval from the Institute’s Ethics Committee (IEC No. INT/IEC/2023/SP/150). Patients aged between three to sixty years with a confirmed diagnosis of Moyamoya disease based on magnetic resonance angiography (MRA) or digital subtraction angiography (DSA) were enrolled in the study. All patients were recruited through the Department of Neurosurgery, PGIMER, Chandigarh, after obtaining written informed consent from the patients or their next of kin. A total of 15 MMD patients were included in the study and peripheral blood samples were collected from each patient. Additionally, patients with acute infarction or haemorrhage (other than MMD) within the past three months were excluded, as well as those with MRI findings indicating a significant area of infarction, haemorrhage, or encephalomalacia. Serum samples from 15 healthy volunteers without any known comorbidities were used as controls.

A total of 5 mL venous blood was collected from each participant in plain red-top vacutainers and allowed to clot at room temperature for 30 minutes. Samples were then centrifuged at 1500× g for 10 minutes, after which the supernatant (serum) was carefully collected. The serum aliquots were snap-frozen at -80 °C until further use.

### Extraction and characterization of serum derived exosomes (SDEs)

SDEs were isolated using pooled serum samples from MMD patients and healthy controls using ultracentrifugation method (Optima XPN 100 ultracentrifuge Beckmann coulter) as shown in our previous study [18]. Serum samples from five individuals were pooled in equal volumes to generate a total volume of 5 mL per pool. Three such pools were prepared from 15 patient samples, and a parallel set of three pools was created from healthy control samples. Each pooled serum sample was subjected to sequential centrifugation at 500×g for 10 minutes and 2000×g for 20 minutes at 4 °C to remove cells and debris. The resulting supernatant was diluted 1:1 with cold 1× phosphate-buffered saline (PBS) and centrifuged at 10,000×g for 30 minutes at 4 °C. The supernatant was then filtered through a 0.22 μm filter and ultracentrifuged twice at 100,000×g for 70 minutes at 4 °C to pellet exosomes. The final exosome pellets were resuspended in PBS and stored at -80 °C until further analysis. SDEs were characterized using electron microscopy, particle size analysis via dynamic light scattering, and confirmed by detecting exosomal markers TSG101, ALIX, LAMP2, and CD63, alongside the non-exosomal marker albumin (ALB) as in our previous study (data not shown here) [18].

### Protein extraction and quantification of SDEs

These SDEs were lysed with Radio Immunoprecipitation Assay (RIPA) lysis buffer and the protein concentration was determined using BCA assay (Pierce™ BCA Protein Assay Kit A65453).

### SDEs proteomic analysis

A total of 50 µg of protein was dissolved in 50 mM ammonium bicarbonate and reduced with 10 µL of dithiothreitol (DTT) at 60 °C for 60 minutes. Subsequently, proteins were alkylated with 10 µL of 20 mM iodoacetamide (IAA) for 30 minutes at room temperature in the dark. The reduced and alkylated protein samples were then digested overnight at 37°C using 1 µg of sequencing-grade trypsin (Promega, #V5111). The digestion was quenched by adding 5 µL of 0.1% formic acid, and peptides were desalted using reversed-phase C-18 spin columns (Thermo Fisher Scientific, #89870). Peptides were eluted in three sequential steps using 30 µL, 40 µL, and 30 µL of elution buffer containing 70% acetonitrile (ACN) and 30% LC-MS grade water. The combined eluates were then concentrated using a speed vacuum concentrator at 45 °C. The Samples were further reconstituted in 0.1% formic acid and were analyzed using LC-MS/MS on a Q-Exactive™ Plus on a Q-Exactive™ Plus instrument (Thermo Fisher Scientific). Data were processed using Proteome Discoverer version 2.0 (Thermo Fisher Scientific), and protein identification was performed against the Uniprot Homo sapiens database (UP000005640) using the Mascot algorithm (Mascot 2.4, Matrix Science). Identified proteins were subjected to statistical, network, and pathway analyses (Fig. 1).

**Fig. 1.**
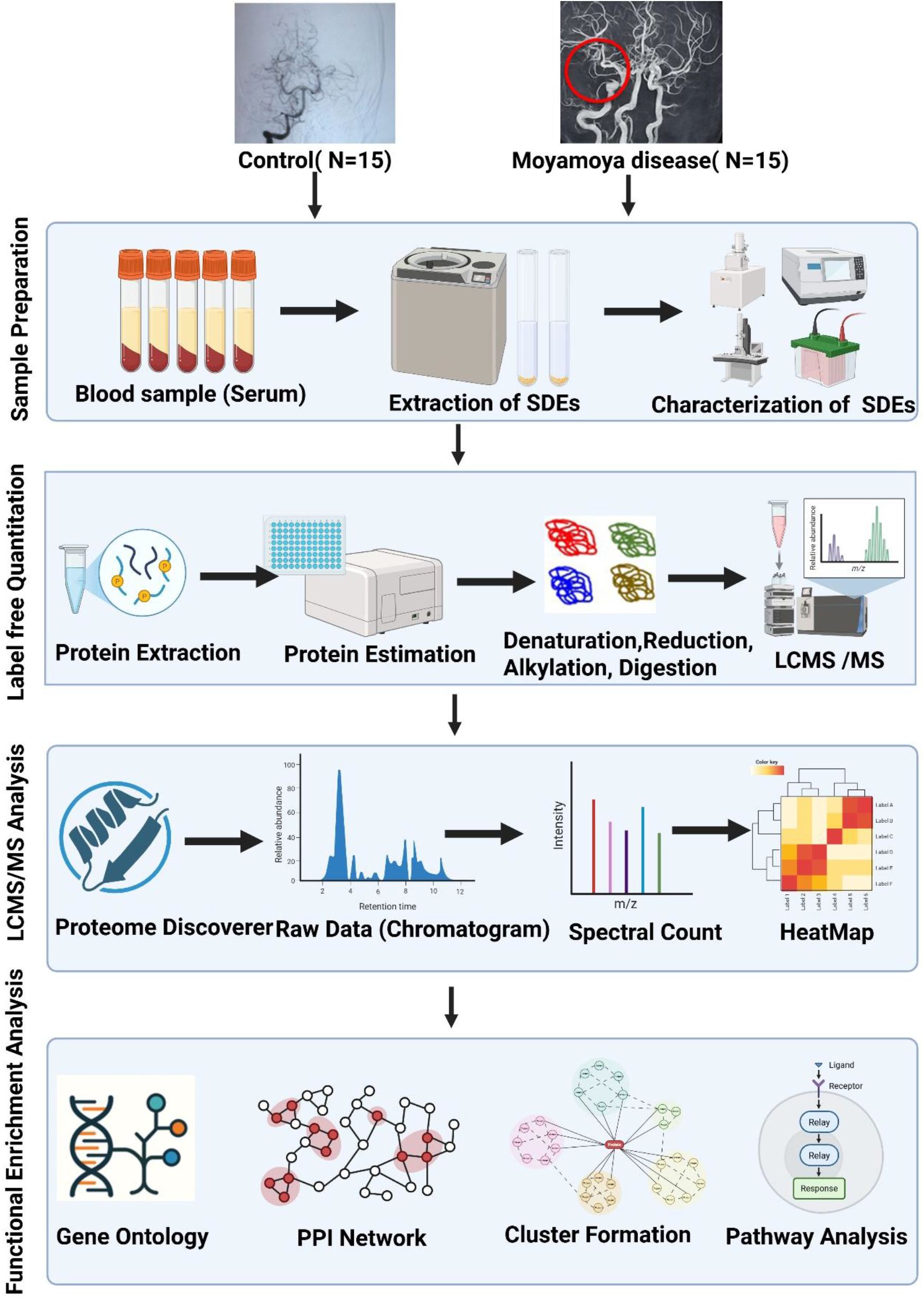
Schematic overview of the experimental workflow for the isolation and proteomic analysis of serum-derived exosomal proteins.

### Bioinformatic analysis

We performed Principal Component Analysis (PCA) using MetaboAnalyst 5.0 (https://www.metaboanalyst.ca) to evaluate the variation in proteomic profiles between the control and Moyamoya Disease groups. Subsequently, a volcano plot was generated to identify significantly altered proteins based on fold change and p-value. Lastly, a heatmap was constructed to visualize protein expression patterns across samples through colour gradients and hierarchical clustering. All analyses were carried out using the MetaboAnalyst platform. Gene Ontology enrichment analysis of dysregulated proteins were performed using FunRich (version 3.1.3) (https://www.FunRich.org), DAVID (https://davidbioinformatics.nih.gov/), and Enrichr tools (https://maayanlab.cloud/enrichr/) [19, 20].

Furthermore, pathway analysis was performed using DAVID (Database for Annotation, Visualization, and Integrated Discovery) [21]. To investigate the functional roles of dysregulated proteins and construct their protein–protein interaction (PPI) network, the STRING database (Search Tool for the Retrieval of Interacting Genes/Proteins, version 11.0) was utilized [22]. A high-confidence score threshold (> 0.9) was applied to construct the protein-protein interaction (PPI) network. The resulting network was exported to Cytoscape (version 3.6.0) for visualization [23], and cluster analysis was performed using the MCODE plugin to identify densely interconnected protein modules. Clusters with a higher number of nodes were selected for further analysis to identify key functional modules. Protein class classification was performed using PANTHER (version 14), a web-based tool for protein annotation and classification [24]. Gene Ontology (GO) terms and pathways with a p-value ≤ 0.05 were considered statistically significant for enrichment analysis.

### Validation of candidate biomarkers

Validation was carried out at both transcript and protein levels using RT-qPCR and ELISA, respectively, in an independent cohort comprising 15 confirmed MMD patients and 15 control samples.

### RT-qPCR

For RT-qPCR, RNA was extracted from serum-derived exosomes using the TRIzol LS method. Briefly, phase separation was carried out with TRIzol LS (Sigma-Aldrich, Cat. No. T3934) reagent and chloroform, followed by RNA precipitation with isopropanol and washing with 75% ethanol. RNA quality was assessed by 1.2% agarose gel electrophoresis, and concentrations were measured using a NanoDrop spectrophotometer (BioTek EPOCH2C). A total of 100 ng RNA samples was used for cDNA synthesis with a High-Capacity cDNA Reverse Transcription Kit (Applied Biosystems, Thermo Fisher Scientific, Cat. No. 4368814). Relative gene expression was then quantified by RT-qPCR using gene-specific primers designed with the NCBI Primer-BLAST tool. The primer sequences used were as follows: RHOA: Forward 5′-GAGCCGGTGAAACCTGAAGA-3′, Reverse 5′-CAAGAAAGTTGGGCACAAGACA-3′; PRKG-2: Forward 5′-TGGAAAGTTACCCGGCTGTC-3′, Reverse 5′-AGCTCTGTCACCTTGTTCCG-3′; MYC: Forward 5′-CCACCTCCAGCTTGTACCTG-3′, Reverse 5′-GAGCAGAGAATCCGAGGACG-3′; GAPDH: Forward 5′-TGAACGGGAAGCTCACTGG-3′, Reverse 5′-TCCACCACCCTGTTGCTGTA-3′. RT-qPCR analysis was performed using DyNAmo ColorFlash SYBR Green Master Mix (2X) (Thermo Fisher Scientific, Cat. No. F416L). Reactions were run in triplicates on a StepOne Real-Time PCR system (Applied Biosystems) with the following program: 95°C for 10 min, followed by 40 cycles of 95°C for 15s, 64°C for 30s, and 72°C for 30s, with a final extension at 72°C for 10 min. Melt curve analysis was performed, and gene expression was normalized to GAPDH.

### Enzyme linked immunosorbent assay (ELISA)

For validation at protein-level, serum samples were analyzed using pre-coated ELISA kits specific for PRKG-2 (ELK Cat. No.-ELK5039), MYC (Finetest Cat. No.- EH2847) and RHOA (Finetest Cat. No.- EH2484) All assays were carried out strictly following the manufacturer’s instructions.

### Statistical Analysis

The statistical data was analysed using GraphPad Prism software version 8.01 (GraphPad Software, San Diego, CA, USA) and Microsoft Excel 2019. An unpaired, Mann-Whitney test (non-parametric test) was used for assessing the statistical significance between groups. A p-value of ˂ 0.05 was considered to be statistically significant. ROC curve analysis was done using GraphPad Prism software. All the experiments were performed in triplicates and data were presented as mean ± standard deviation.

## Results

### Demographic details of MMD patients and controls

In the discovery phase, the patient cohort comprised paediatric cases (n = 10; mean age 7.6 ± 4.06 years) and adult cases (n = 5; mean age 43.6 ± 12.23 years), reflecting the bimodal age distribution of MMD. The female-to-male ratio among patients was 8:7, with the majority presenting with ischemic symptoms. In the control group, the female-to-male ratio was 7:8.

### Functional and pathway analysis

Principal Component Analysis (PLS-DA) was conducted using MetaboAnalyst 5.0 software to assess variation in proteomic profiles between the control and Moyamoya Disease groups. The resulting scores plot demonstrates a clear separation between the two groups, indicating distinct protein expression patterns (Fig. 2). The first principal component (PC1) captures 26.7% of the total variance, while the second component (PC2) accounts for 21.4%, together representing a substantial portion of the overall variability in the dataset. Differential expression analysis was conducted on 2,554 proteins, revealing 213 proteins as significantly dysregulated (p ≤ 0.05) with a Log_2_ fold change ≥ 2 or ≤ -2. Among these, 118 proteins were markedly upregulated (Log_2_ fold change ≥ 2), while 95 proteins were significantly downregulated (Log_2_ fold change ≤ -2). The distribution of differentially expressed proteins is illustrated in the volcano plot and heatmap generated using MetaboAnalyst software 5.0 (https://www.metaboanalyst.ca).

**Fig. 2.**
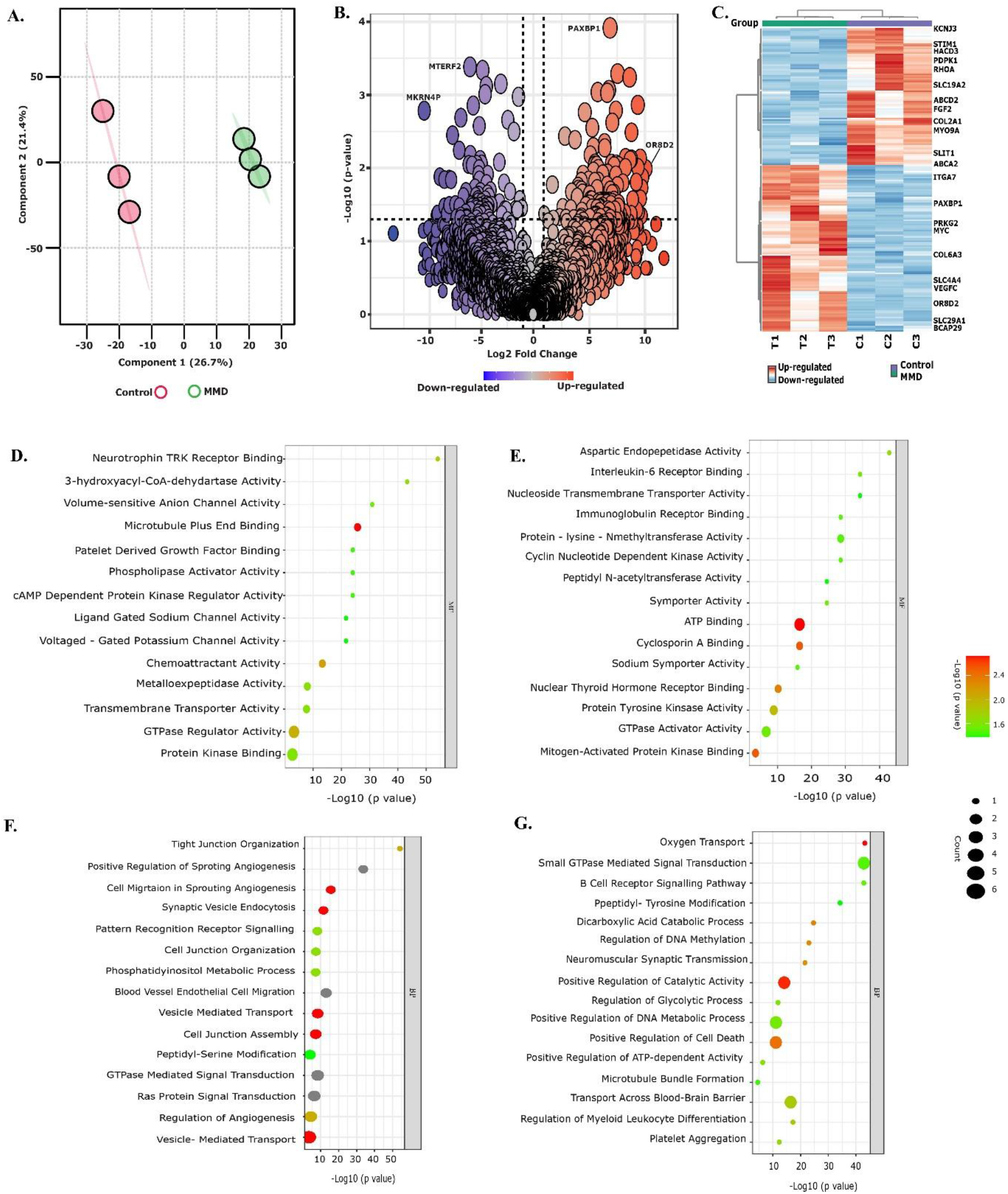
**(A)** PCA of proteomic profiles from MMD patients and controls shows clear group separation, with PC1 and PC2 capturing the major variance, indicating distinct protein expression patterns between groups. **(B)** Volcano plot illustrates differentially expressed proteins: red (upregulated), blue (downregulated), and black (non-significant) in MMD versus controls. **(C)** Unsupervised hierarchical clustering heatmap displays significantly dysregulated proteins, with clear separation between MMD (green) and controls (blue), highlighting consistent expression differences. **(D & E)** Bubble plots illustrating enriched Molecular Functions (MF) Gene Ontology (GO) for downregulated and upregulated proteins, respectively. **(F & G)** Bubble plots of Biological Processes (BP) GO for downregulated and upregulated proteins, respectively. Bubble colour indicates statistical significance whereby, red color indicates higher log10 (pvalue) and green color indicate lower log10 (pvalue) and size of the bubble reflects the number of associated proteins.

### Functional landscape of the serum-derived exosomal proteome in Moyamoya disease

Using Gene Ontology (GO) analysis, we categorized the dysregulated proteins identified in Moyamoya disease according to their associated molecular functions and biological processes.

### Molecular function

Functional enrichment analysis of the upregulated proteins demonstrated significant enrichment in key molecular activities, particularly the downregulated proteins were primarily enriched in molecular functions such as neurotrophins TRK receptor binding (PLCG1), microtubule plus-end binding (DST, STIM1), platelet-derived growth factor binding (COL2A1), phospholipase activator activity (PDPK1), and chemoattractant activity (COLEC10, FGF2) (Fig. 2D). In contrast, the upregulated proteins particularly ATP binding (ABCA2, RTEL1, TNK1, ABCB8, MORC2, EIF4G1), protein tyrosine kinase activity (BLK, CLK1, TNK1), GTPase activator activity (RANBP2, ARHGAP11B, ARHGAP40, TAGAP), and MAP kinase binding (PRKG2, ATF7). These functional categories are closely associated with intracellular signalling, metabolic activity, and cytoskeletal remodeling (Fig. 2E). Additionally, a significant number of downregulated proteins were involved in GTPase regulator activity (DENND1C, ARHGEF37, TNK2, RapGEF1, HACD3, MYO9A), and protein kinase binding (PPP1CB). Notably, several of these proteins such as ARHGEF37, MYO9A, and RapGEF1 are upstream modulators of RHOA signalling, implicating a disruption in RHOA-mediated cytoskeletal regulation and junctional dynamics in the downregulated protein network.

### Biological process

The biological process enrichment analysis further highlighted distinct functional signatures between upregulated and downregulated proteins. Downregulated proteins were enriched in processes including: tight junction organization (PATJ), positive regulation of sprouting angiogenesis (PDPK1, FGF2), regulation of cell migration during sprouting angiogenesis (FGF2, RHOA), endothelial cell migration (PDPK1, PLCG1, FGF2), cell junction organization and assembly (DNM3, MYO9A, RHOA), overall regulation of angiogenesis (STIM1, PDPK1, PLCG1, FGF2). These downregulated processes highlight impairments in endothelial barrier integrity, angiogenic sprouting, and directed migration (Fig. 2F). The repression of RHOA and its regulatory partners (ARHGEF37, RAPGEF1, MYO9A) indicates disrupted RHOA GTPase signalling, which is essential for cytoskeletal rearrangement, junctional complex assembly, and vascular morphogenesis. Upregulated proteins on the other hand were significantly involved in: oxygen transport (HBB, HBA1), positive regulation of extracellular matrix (ECM) disassembly (CLASP1), regulation of heme biosynthesis (SLC6A9), transport across the blood-brain barrier (SLC6A9, SLC29A1, SLC4A4), platelet aggregation (UBASH3B, HBB), positive regulation of cell death (HBB, HBA1, EIF4G1), microtubule bundle formation (NCKAP5, CLASP1). These findings suggest that the upregulated protein profile supports enhanced tissue oxygenation, matrix remodeling, and nutrient exchange processes critical for endothelial activation and angiogenesis (Fig. 2G). Furthermore, increased platelet aggregation and regulated cell death may contribute to pro-angiogenic signalling, while microtubule organization is likely essential for endothelial migration and vessel stabilization.

### Pathway enrichment analysis of dysregulated proteins

Pathway enrichment analysis revealed significant upregulation of several signalling cascades, notably focal adhesion, PI3K-Akt, axon guidance, neurotrophin signalling, ABC transporters, and NF-κB signalling. Among these, the focal adhesion and PI3K-Akt pathways were most prominently enriched and are intimately linked to the regulation of cytoskeletal dynamics, cell proliferation, and survival, key processes central to vascular smooth muscle cell (VSMC) phenotypic switching (Fig. 3A). Within the focal adhesion pathway (Fig. 3B), multiple components were dysregulated, including extracellular matrix (ECM) proteins (COL2A1, COL6A3), growth factors (VEGFC, FGF2), integrins (ITGA7), and regulators of cytoskeletal reorganization such as the small GTPase RHOA and its modulators (RhoGEFs and RhoGAPs). Downstream effectors like myosin light chain phosphatase (MLCP) and PDPK1, a kinase that activates Akt, also showed altered expression. (Fig. 4A shows a bubble plot representing all enriched pathways identified using the SR plot).

**Fig. 3.**
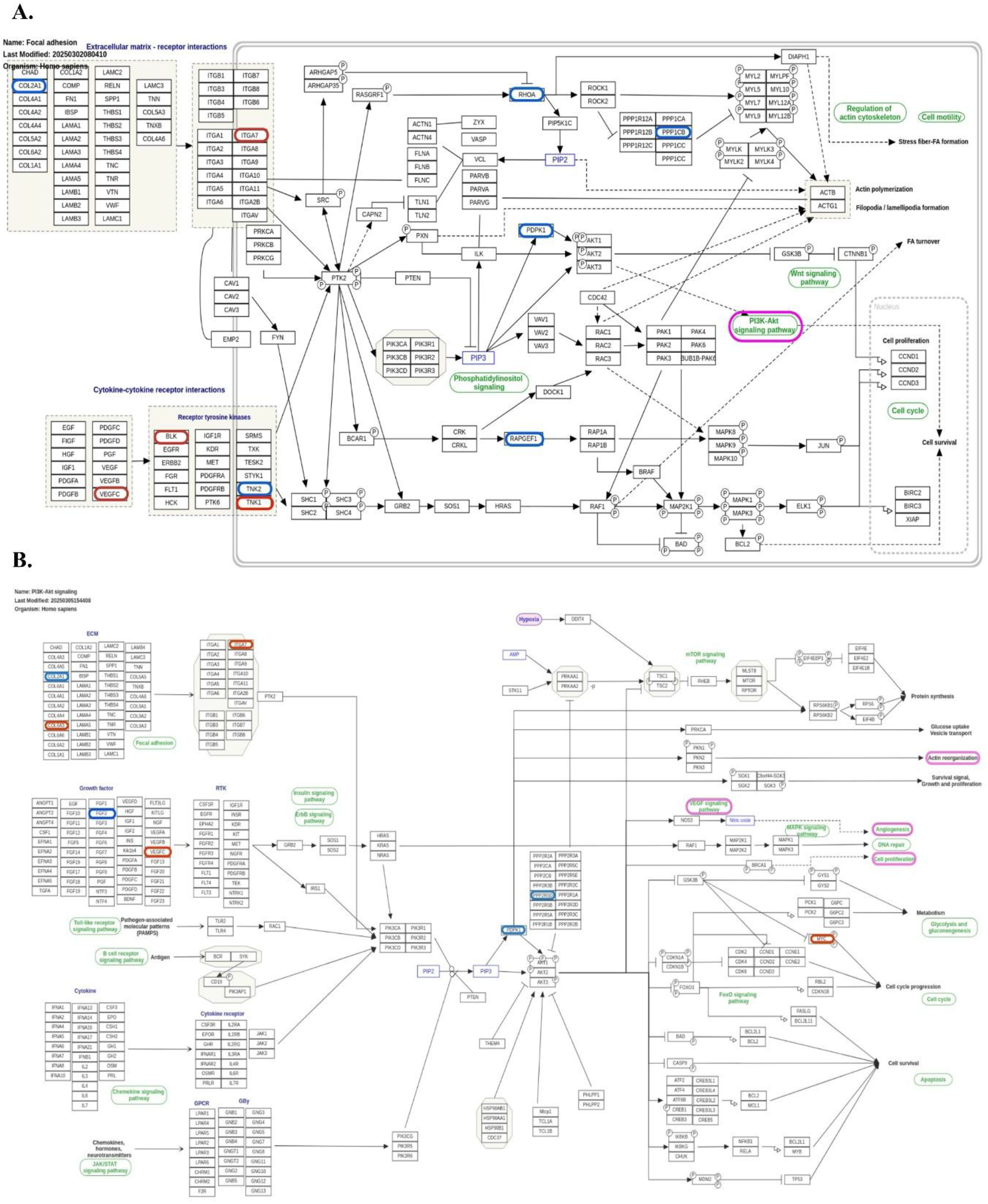
**(A)** Schematic illustration of the focal adhesion signalling pathway depicting key protein– protein interactions. (**B)** Schematic illustration of the PI3-AKT signaling pathway. Differentially expressed or pathway-relevant proteins are outlined in color-coded borders: blue boxes represent downregulated proteins, while red boxes denote upregulated proteins.

**Fig. 4.**
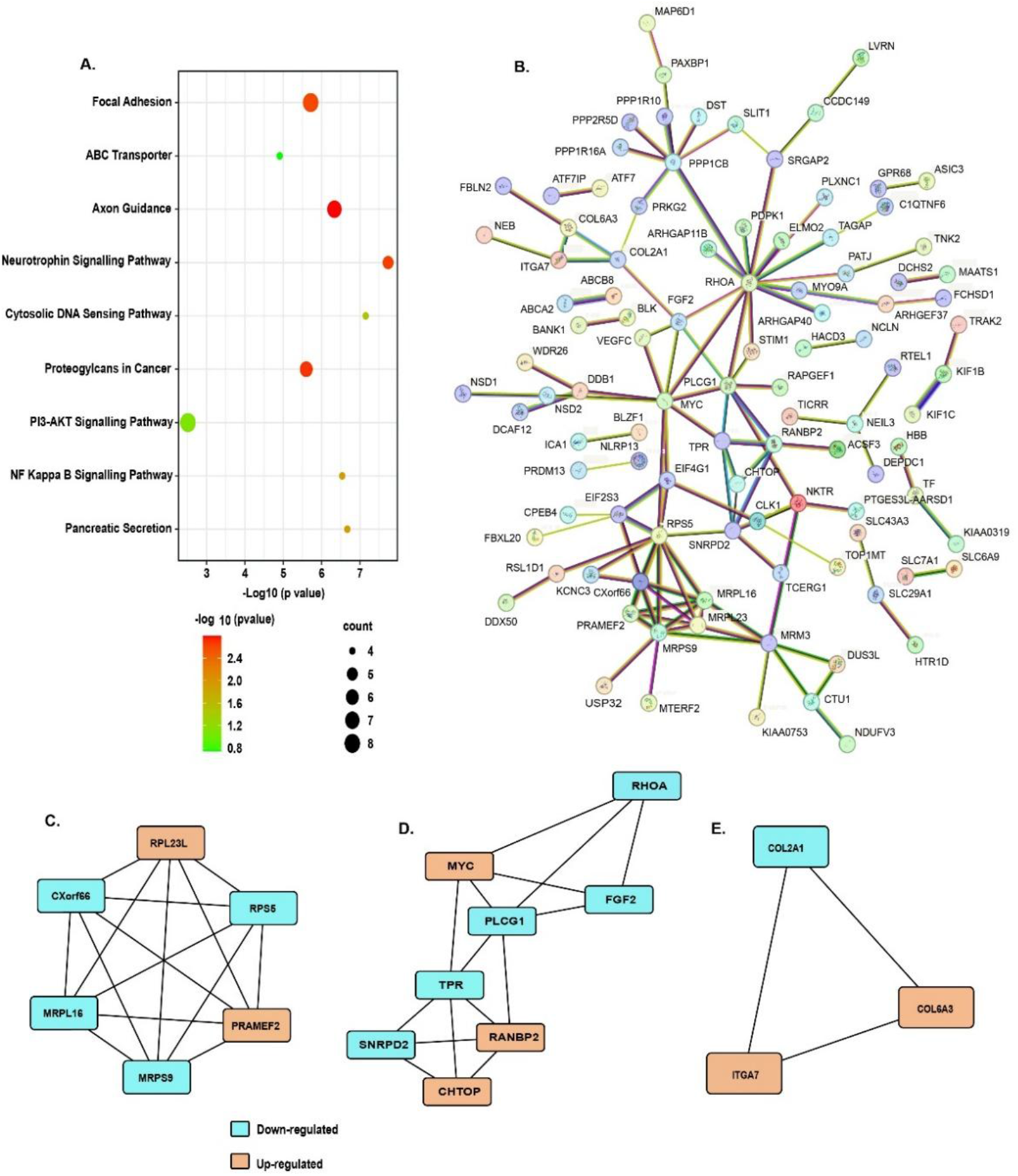
**(A)** KEGG pathway enrichment analysis showing significantly enriched pathways. **(B)** Protein– protein interaction (PPI) network constructed using STRING database for the identified differentially expressed proteins. Nodes represent proteins, and edges represent known or predicted associations. **(C-E)** Clusters of differentially expressed proteins.

The focal adhesion and PI3K-Akt pathways appear to be interconnected at multiple nodes, as both are activated by common ligands such as ECM proteins and growth factors, and by receptors including integrins and tyrosine kinases. Furthermore, these pathways converge at PDPK1, a key molecular junction and upstream activator of Akt, whose expression was found to be markedly downregulated. Additional dysregulated molecules such as PPP1R16A and PPP2R5D, known inhibitors of Akt signalling, were also identified, suggesting altered regulatory control. Notably, MYC, a downstream transcription factor in the Akt-GSK3 signalling axis, was found to be upregulated, indicating enhanced proliferative signalling.

Although less enriched, other identified pathways were also associated with critical biological functions such as neural protection, intercellular communication, and immune responses, highlighting a broader landscape of signalling alterations in Moyamoya disease.

### Protein-Protein interaction (PPI) network analysis

Protein-protein interaction (PPI) network analysis of 213 dysregulated serum-derived exosomal proteins using the STRING database generated a network consisting of 199 nodes and 123 edges, suggesting moderate interaction density (Fig. 4B). Several hub proteins, including RHOA, FGF2, and HBB, emerged as central regulators with high connectivity. Cluster analysis revealed distinct functional modules within the network. A major cluster cantered around RHOA, FGF2, PLCG1, and PDPK1 comprised proteins involved in cytoskeletal remodeling, angiogenesis, and cell migration, highlighting disruption in RHOA signalling and endothelial dyna22mics in MMD. Another prominent cluster involving HBB, HBA1, SLC6A9, and SLC29A1 was enriched for molecules linked to oxygen transport, platelet aggregation, and blood-brain barrier regulation, indicating compensatory responses to chronic cerebral hypoperfusion. A third cluster, including ITGA7, COL2A1, and COL6A3, reflected changes in extracellular matrix-receptor interactions and focal adhesion signalling, supporting active vessel wall remodeling. Additionally, a cluster containing MYO9A, ARHGEF37, and RapGEF1 represented key regulators of RhoGTPase activity and tight junction organization, suggesting impaired endothelial barrier function and vascular smooth muscle cell (VSMC) phenotypic switching. Overall, the network underscores a coordinated dysregulation of pathways central to vascular remodeling, cytoskeletal regulation, and metabolic adaptation, offering valuable insight into the molecular mechanisms driving MMD pathogenesis.

Further, using Cytoscape, protein-protein interaction (PPI) networks were also constructed to identify functionally relevant gene clusters among the differentially expressed proteins. Three prominent protein modules were identified: (i) Mitochondrial and Ribosomal Protein-Enriched Module, (ii) Transcriptional Regulation and Signal Transduction Hub, and (iii) Extracellular Matrix Remodeling and Adhesion Complex. The mitochondrial and ribosomal module includes proteins such as RPL23L, MRPL16, MRPS9, RPS6, and PRAMEF2, with most being down-regulated, indicating a potential suppression of mitochondrial translation and energy metabolism (Fig. 4C). The up-regulation of PRAMEF2 may reflect a compensatory or stress-responsive mechanism. The transcriptional and signalling network involves a complex network cantered on transcriptional regulators and signalling mediators such as MYC, PLCG1, FGF2, and RHOA (Fig. 4D). This module includes both up-regulated (MYC, RANBP2, CHTOP) and down-regulated (FGF2, RHOA) nodes, suggesting a dynamic balance between proliferative signalling and regulatory feedback loops. This cluster may reflect the dysregulation of pathways involved in cell cycle control, transcriptional regulation, and cellular signalling, potentially driving pathophysiological remodeling or disease progression. The extracellular matrix-related module comprises COL2A1 (down-regulated) along with COL6A3 and ITGA7 (up-regulated) (Fig. 4E), indicating active ECM remodeling and altered cell-matrix adhesion, which may underlie structural tissue changes or enhanced cellular invasiveness.

### Validation of candidate biomarkers in serum-derived exosomes

Candidate molecules, including PRKG-2, MYC, and RHOA, were validated at both transcript and protein levels in serum-derived exosomes. Validation was performed using serum samples from an independent cohort comprising Moyamoya disease patients (n = 15; mean age 21.5 ± 16.62 years; female-to-male ratio 9:6) and matched controls (female-to-male ratio 5:10). Among the MMD samples, 7 were from adult patients and 8 from paediatric patients, whereas in the control group, 9 were adults and 6 were paediatric subjects. PRKG2 and MYC were significantly upregulated in MMD patients compared with controls, whereas RHOA was markedly downregulated at both the transcript and protein levels in MMD patients. At the transcript level, PRKG2 expression was increased by ∼4.99 fold (p < 0.0001) (Fig. 5A) and MYC by ∼1.35 fold (p < 0.0001) in MMD patients (Fig. 5D), whereas RHOA showed a significant reduction with a log₂-fold change of -1.72 (p < 0.0001) (Fig. 5G).

**Fig. 5.**
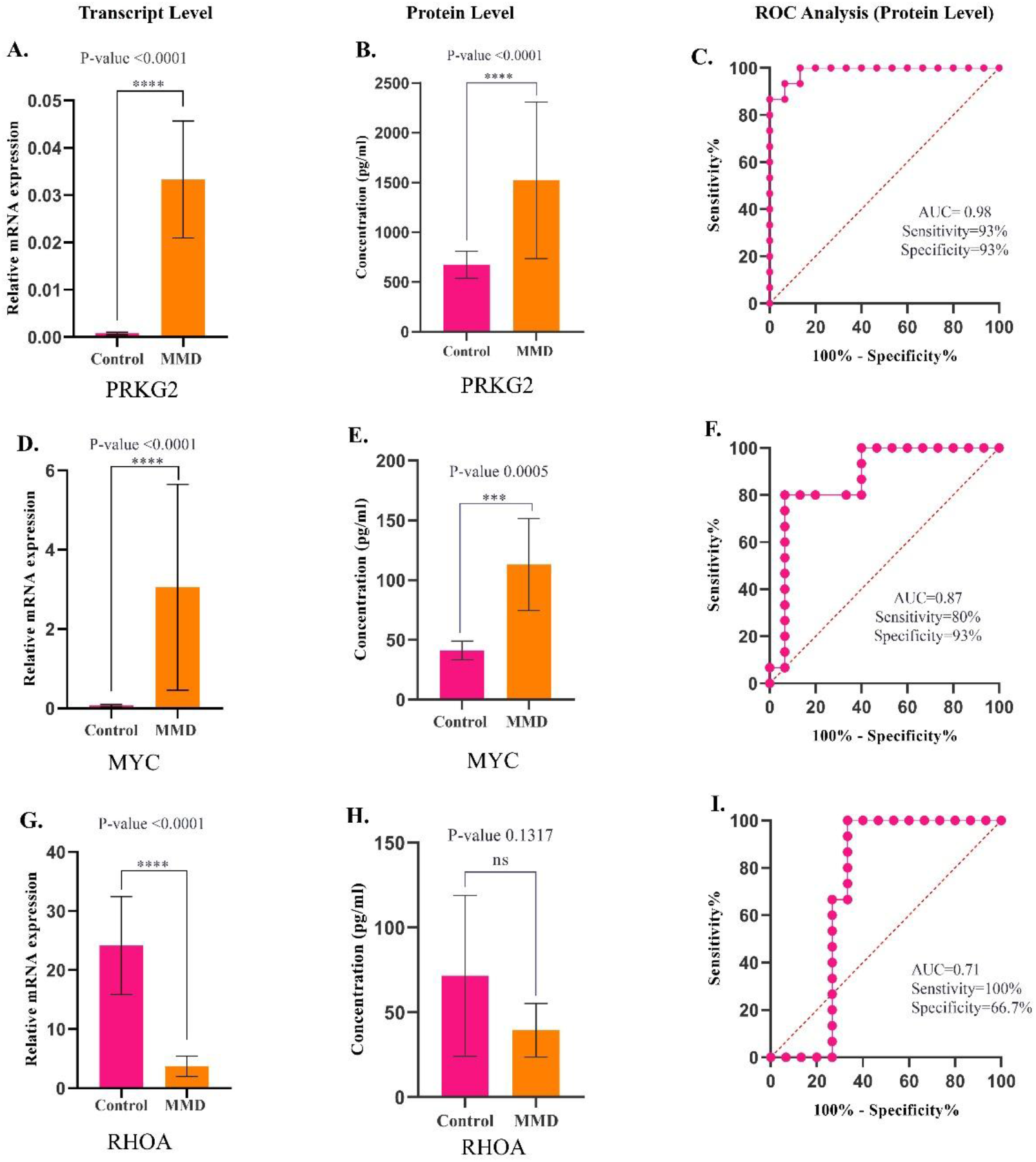
Validation of candidate proteins at the transcript and protein levels using qRT-PCR and ELISA, respectively. Panels **(A)**, **(D)**, and **(G)** represent the relative mRNA expression levels of PRKG2, MYC, and RHOA, respectively. Panels **(B)**, **(E)**, and **(H)** depict the corresponding protein expression levels. Receiver Operating Characteristic (ROC) curve analyses for PRKG2, MYC, and RHOA are shown in panels **(C)**, **(F)**, and **(I)** respectively.

At the protein level, PRKG2 and MYC were significantly upregulated, showing a log₂-fold increase of 10.58 (P < 0.0001) (Fig. 5B) and 7.17 (P = 0.0005), respectively, compared to controls (Fig. 5E). In contrast, RHOA was downregulated with a log₂-fold change of 5.31 (P = 0.1317), which was not statistically significant (Fig. 5H).

Receiver Operating Characteristic (ROC) curve analysis revealed that PRKG2 exhibited excellent diagnostic performance with an AUC of 0.98, 93% sensitivity, and 93% specificity (Fig. 5C), whereas MYC showed strong discriminative ability with an AUC of 0.87, 80% sensitivity, and 93% specificity (Fig. 5F). In contrast, RHOA displayed moderate discrimination with an AUC of 0.71, 100% sensitivity, and 66.7% specificity (Fig. 5I).

## Discussion

In light of the limited understanding of the molecular mechanisms underlying Moyamoya disease, growing attention has shifted to exosomes with emerging diagnostic and therapeutic potential. Secreted by nearly all cell types and present in various bodily fluids, exosomes reflect the physiological or pathological state of their cells of origin. Emerging *in vitro* and *in vivo* evidence suggests that exosomes may play a role in the pathophysiology of cerebrovascular disorders, including MMD. To investigate the molecular underpinnings of MMD, we conducted proteomic profiling of serum-derived exosomes (SDEs) using LC-MS/MS, revealing substantial differences between MMD and control groups. Out of 2,554 proteins identified, 213 were significantly dysregulated (Log₂ FC ≥ 2 or ≤ -2, p ≤ 0.05), comprising 118 upregulated and 95 downregulated proteins. This molecular divergence suggests systemic alterations in signalling, inflammation, vascular remodeling, and neurovascular regulation in MMD.

Further, gene ontology (GO) analysis of dysregulated proteins in MMD revealed altered molecular functions including protein tyrosine kinase activity, GTPase activator activity, MAP kinase binding, and phospholipase activator activity, indicating disruptions in signalling, metabolism, and cytoskeletal regulation. These are critical for endothelial function and vascular homeostasis. Biological process analysis showed involvement in endothelial migration, angiogenesis, ECM disassembly, tight junction organization, and microtubule dynamics which are key components of vascular remodeling. Upregulation of proteins related to ECM breakdown and cytoskeletal changes suggests compensatory angiogenic responses to chronic hypoperfusion. In contrast, downregulated proteins affecting junction integrity and pro-angiogenic signalling imply impaired vascular repair and endothelial plasticity. Additionally, the dysregulated expression of VEGF, LY6G6F, integrins, and collagen in our dataset reflects active VSMC phenotypic switching, characterized by loss of contractile features and acquisition of synthetic, proliferative, and migratory traits. Such transitions facilitate VSMC intimal migration and ECM remodeling, thereby reinforcing fibro-cellular thickening and maladaptive vascular remodeling. Collectively, these molecular alterations point to dysregulated pathways, including RHOA, PI3K-Akt, and focal adhesion that drive maladaptive vascular remodeling and may contribute to vessel stenosis and collateral formation in Moyamoya disease.

### Mechanisms of vascular remodeling and angiogenic response in MMD

#### I. Vascular occlusion

Intimal thickening lies at the core of vascular remodeling in Moyamoya disease [25]. It involves pathological thickening of the vessel wall, primarily driven by vascular smooth muscle cell (VSMC) proliferation and migration to intima. This is often initiated by endothelial dysfunction and altered local signalling [25, 26] (Fig. 6).

**Fig. 6.**
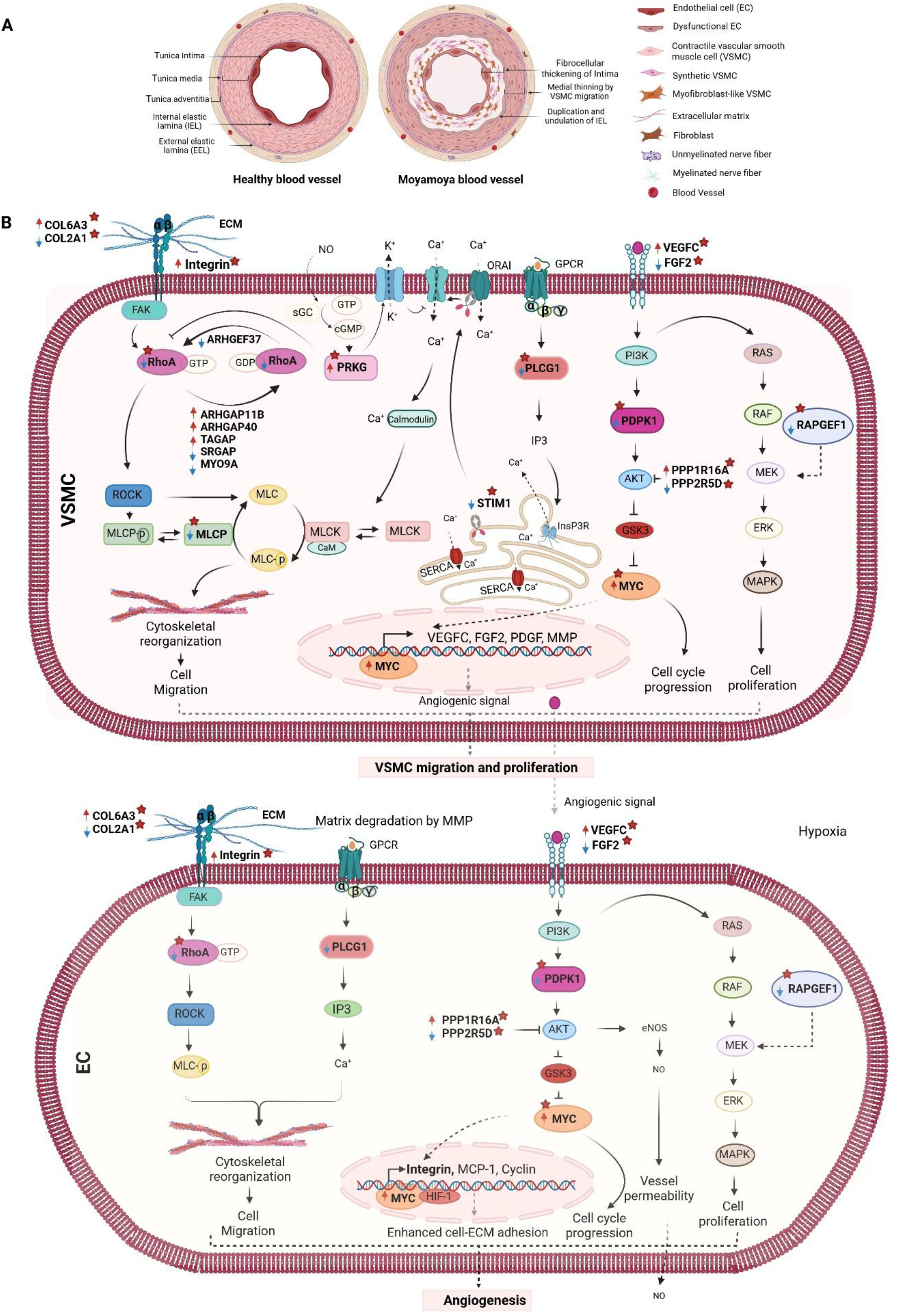
**Vessel wall remodeling in Moyamoya disease. (A) Comparison of a healthy vessel (left) with a Moyamoya vessel (right).** In Moyamoya disease, endothelial dysfunction and aberrant growth factor signalling promote vascular smooth muscle cell (VSMC) dedifferentiation, proliferation, and migration from the media into the intima. This results in fibro-cellular thickening, medial thinning, and duplication/undulation of the internal elastic lamina (IEL). Together with extracellular matrix (ECM) deposition and VSMC phenotypic transitions, these changes drive intimal hyperplasia and progressive vascular occlusion. **(B) Signalling mechanisms driving VSMC migration/proliferation and EC-mediated angiogenesis in Moyamoya disease.** Growth factors such as VEGF and FGF, along with ECM remodeling, activate integrin-, GPCR-, and PI3K-Akt-dependent pathways in both vascular smooth muscle cells (VSMCs) and endothelial cells (ECs). In VSMCs, these cues promote phenotypic switching, cytoskeletal reorganization through RHOA/ROCK/MLC and Ca²⁺ signalling, and activation of MAPK/MYC pathways, leading to migration, proliferation, and intimal hyperplasia. In parallel, ECs respond to hypoxia and VSMC-derived growth factors by initiating cytoskeletal remodeling, Ca²⁺ influx, and eNOS/NO signalling. Importantly, NO released from ECs activates PRKG in VSMCs, which suppresses RHOA activity and modulates Ca²⁺-dependent cytoskeletal dynamics, thereby fine-tuning VSMC motility. Conversely, VSMCs secrete angiogenic factors (VEGF, FGF, PDGF, MMPs) that stimulate EC proliferation, migration, and sprouting. This reciprocal signalling establishes a feed-forward loop in which VSMC-driven intimal hyperplasia and EC-driven angiogenesis act in concert to drive progressive vascular remodeling in Moyamoya disease.

Endothelial-derived growth factors can activate intracellular pathways that induce VSMC phenotypic modulation, promoting their transition from a contractile to a synthetic state [27]. Our proteomic analysis of serum-derived exosomes from MMD patients supports these mechanisms, revealing upregulation of vascular endothelial growth factor (VEGF), likely reflecting a hypoxia-induced angiogenic response. Elevated expression of LY6G6F, a member of the Ly6a stem cell marker family, may reflect vascular smooth muscle cell phenotypic modulation into mesenchymal type. This is also substantiated by the down regulation of contractile proteins. Concurrent changes in ECM proteins such as COL6A3 and COL2A1 (collagen), along with ITGA7 (integrin), suggests active-matrix remodeling and enhanced integrin-ECM interactions facilitating VSMC migration.

VSMCs exhibit remarkable plasticity, enabling them to transition from a quiescent, contractile phenotype to synthetic or mesenchymal-like states. This phenotypic modulation is characterized by downregulation of contractile markers and acquisition of proliferative, migratory, and ECM-secreting traits [28]. Synthetic VSMCs typically express ECM-related and PDGF-responsive markers such as FN1, COL1A1, and PDGFRβ, whereas progression toward mesenchymal-like or progenitor-like states is indicated by the induction of Ly6 family proteins, reflecting deeper dedifferentiation and enhanced cellular plasticity.

### Mediators of VSMC migration in disease: from the vessel wall to the intima

#### Growth factors

Growth factors are central regulators of VSMC phenotypic modulation. PDGF-BB is a well-established driver of this switch, acting through multiple mechanisms: upregulating the transcription factor KLF4, suppressing myocardin (MYOCD), and disrupting its binding to contractile gene promoters [28, 31]. VEGF also plays a direct role in VSMC phenotypic switching via STAT3 and ERK1/2 signalling [32, 33]. Our proteomic analysis revealed elevated levels of VEGF and reduced levels of FGF2 in serum-derived exosomes from MMD patients.

#### Integrins

Integrins, key transducers of ECM signals, facilitate VSMC adhesion, migration, and proliferation via cytoskeletal remodeling and focal adhesion turnover [34]. Integrins interact with collagen, and their expression is upregulated by growth factors and mitogens [35]. Our study showed upregulation of ITGA7 and COL6A3 (collagen) in MMD serum, indicating enhanced integrin-collagen engagement and activation of focal adhesion pathways.

#### Microtubules

Microtubules orchestrate cell polarity and directional migration by working with actin filaments and integrins [36]. During phenotypic switching, they guide the formation of lamellipodia and stress fibres [37]. In MMD, dysregulation of microtubule-related proteins, including upregulated MAP6D1 and CYCL1, and downregulated MAST3, RHOA1, ODF3B, and PATJ, suggests impaired cytoskeletal coordination and disrupted cell motility.

#### Cytoskeletal reorganization

Cytoskeletal remodeling is a key driver of VSMC contraction, migration, and phenotypic switching, core processes in Moyamoya disease pathophysiology [38]. The cytoskeleton, composed of actin filaments, microtubules, and intermediate filaments, is dynamically regulated through focal adhesions linking integrins to actin, activating downstream pathways such as MAPK, PDPK1, and phosphoinositide signalling [39, 40]. Small GTPases, particularly RHOA, RAC1, CDC42, and RAP1, coordinate actin polymerization and myosin-based contractility [41]. RHOA, via its effector ROCK, promotes Ca²⁺-independent activation of myosin II by inhibiting myosin light chain phosphatase (MLCP), thereby promoting VSMC contractile tone [42]. In addition, RHOA regulates cytoskeletal integrity, focal adhesion turnover, and transcriptional programs such as the SRF-myocardin axis to maintain VSMC identity [43].

#### Transcription factors

Transcription factors are central regulators of VSMC phenotypic plasticity and vascular remodeling. Among them, KLF4, together with OCT3/4, SOX2, and MYC, was originally identified as a key inducer of somatic cell pluripotency [44]. In our proteomic data, we observed upregulation of MYC, implicating it in pathological vascular remodeling. MYC is a potent transcription factor with over 25,000 potential genomic binding sites [45, 46]. Beyond its proliferative role, MYC regulates cytoskeletal dynamics by modulating RHOA transcription, a key regulator of actin polymerization, contractility, and migration. While the MYC-RHOA axis can drive migration and transformation, MYC also limits RHOA-induced stress fiber assembly and focal adhesion maturation [47]. This dual action results in cytoskeletal relaxation, reduced contractile gene expression, and a shift toward a synthetic, migratory VSMC phenotype [48]. In MMD, elevated MYC coupled with suppressed RHOA activity, as reflected in our exosomal proteome, likely disrupts cytoskeletal homeostasis and promotes VSMC dedifferentiation, thereby driving maladaptive vascular remodeling.

### II. Angiogenic remodeling in MMD

In response to luminal occlusion caused by VSMC migration and fibro-cellular intimal thickening in the terminal internal carotid artery and its branches, fragile collateral vessels develop at the brain’s base through angiogenesis [2]. This process involves new vessel formation from existing vasculature. Endothelial cell migration plays a pivotal role in this process [49]. It is orchestrated by three primary mechanisms: mechanotaxis, chemotaxis, and haptotaxis [50]. Mechanotaxis, mediated by fluid shear stress, regulates EC polarity, adhesion, and motility [51]. Chemotactic migration is driven by gradients of angiogenic factors such as VEGF, bFGF, and angiopoietins, which activate actin-dependent pathways [52]. Haptotaxis involves integrin-mediated interactions with ECM components, particularly collagen, guiding directional migration independent of soluble cues [53]. These processes converge at RHOA mediated focal adhesion signalling cascade, where integrins link ECM to the actin cytoskeleton and initiate signalling cascades amplified by small GTPases (RHOA, RAC1, CDC42) [54].

#### RHOA-mediated crosstalk in vascular remodeling and angiogenesis in MMD

Cytoskeletal remodeling is central to vascular smooth muscle cell (VSMC) phenotypic switching, migration, and contraction as well as in the endothelial migration and adhesion at the new location in angiogenesis [38]. RHOA regulates cytoskeletal integrity, focal adhesion turnover, and transcriptional programs such as the SRF-myocardin axis to modulate vascular remodelling [43, 55]. Beyond its role in movement, RHOA-ROCK signalling can also promote VEGF gene expression, particularly under low-oxygen conditions [56]. It does so by increasing MYC binding to the VEGF promoter, thereby boosting VEGF production from non-endothelial cells and reinforcing the angiogenic response [57]. This finding underscores the pathological significance of disrupted RHOA signalling in the vascular occlusion and impaired angiogenic response characteristic of Moyamoya disease.

RHOA activity is finely regulated by RhoGEFs (activators) and RhoGAPs (inhibitors). In our study, both RHOA and its upstream activator ARHGEF37 were downregulated in serum-derived exosomes from MMD patients, indicating impaired RHOA-ROCK signalling. This aligns with findings from Koh et al. (2025), who reported that extracellular vesicles (EVs) from MMD patients contain elevated levels of miR-512-3p, which directly targets and suppresses ARHGEF3, another critical RHOA activator, thereby reducing RHOA expression [58]. Importantly, restoration of ARHGEF3, and consequently RHOA, expression in MMD-derived endothelial colony-forming cells (ECFCs) significantly improved tubule formation [58]. Consistent with impaired RHOA activation, we also observed upregulation of several RhoGAPs, including ARHGAP11B, ARHGAP40, TAGAP, and ELMO2, which may further suppress RHOA signalling. Conversely, downregulation of other RhoGAPs such as MYO9A and SRGAP2 points to a context-dependent imbalance rather than uniform suppression. This disruption extended beyond RHOA, with reduced expression of RapGEF1 (essential for endothelial barrier stability [59] and TNK2, a CDC42 effector regulating migration [60], suggesting a broader impairment in small GTPase-mediated cytoskeletal dynamics.

Counterbalancing the RHOA-ROCK cascade, the NO-cGMP-PRKG1 axis emerges as a key regulatory pathway in Moyamoya disease. Protein kinase G (PKG), a serine/threonine kinase, is central to vascular smooth muscle contractility [61]. PRKG1 lowers intracellular Ca²⁺ by inhibiting IP₃R-mediated release and enhancing SERCA (Sarco/Endoplasmic Reticulum Ca²⁺-ATPase) activity [62]. It also activates myosin light chain phosphatase (MLCP) while suppressing RHOA-ROCK signalling, thereby promoting cytoskeletal relaxation and reducing contractile stress [63]. Our proteomic analysis revealed upregulation of PRKG1, accompanied by downregulation of STIM1 (stromal interaction molecule 1) [64] and PLCG1 (phospholipase Cγ1) [65], two pivotal regulators of Ca²⁺-dependent signalling. STIM1, an ER membrane calcium sensor, detects ER Ca²⁺ depletion and activates Orai channels at the plasma membrane to replenish cytosolic Ca²⁺ [66]. Elevated STIM1 expression has previously been reported in proliferative VSMCs, including the neointima of injured rat carotid arteries and in VSMCs stimulated by serum and growth factors [67], highlighting its importance in pathological vascular remodeling.

Beyond its role in calcium sensing, the reduction of PLCG1 in our dataset further indicates disrupted Ca²⁺-dependent signalling in MMD. Normally, activated PLCγ1 hydrolyses PIP₂ into IP₃ and DAG; IP₃ releases Ca²⁺ from the ER, while DAG activates PKC. Together, these messengers promote VSMC proliferation, migration, phenotypic switching, and the cytoskeletal changes required for endothelial cell migration and tubulogenesis [67]. Therefore, reduced PLCG1 likely weakens these Ca²⁺-driven processes, limiting both vascular remodeling and angiogenesis [65]. Importantly, the imbalance between RHOA-ROCK-Ca²⁺-driven contractility and NO-PRKG1 -mediated relaxation creates a pathological disequilibrium. While RHOA downregulation may relieve suppression of NO production [68] and enhance PRKG2 activation. Excessive PRKG2 activity combined with reduced STIM1/PLCG1 expression may blunt VSMC plasticity and weaken EC migration. This dual impairment disrupts coordinated VSMC-EC crosstalk, fostering abnormal vessel wall remodeling and contributing to the formation of fragile, disorganized collaterals characteristic of Moyamoya disease.

#### Disrupted signalling pathways underlying vascular remodeling and angiogenesis in MMD

Pathway enrichment analysis revealed significant involvement of focal adhesion, PI3K-Akt, axon guidance, and proteoglycan-related signalling in Moyamoya disease, indicating a complex disruption of vascular remodeling processes. Dysregulated expression of key proteins such as PDPK1, FGF2, MLCP, PLCG1, and RHOA suggests impaired endothelial cell migration and vascular smooth muscle cell function [69–71]. Focal adhesions, critical for linking integrins to the actin cytoskeleton and enabling VSMC contraction via the RHOA-ROCK-myosin II axis [71], were notably affected in our serum-derived exosomal proteomic data. Similarly, the PI3K-Akt signalling pathway, a central mediator of EC migration and angiogenic survival [72, 73], showed evidence of functional compromise. Reduced PDPK1 expression, together with dysregulation of negative regulators such as PPP2R5D and PPP1R16A, may limit the angiogenic adaptation required under ischemic stress.

Disruption was also evident in axon guidance signalling, a pathway increasingly recognized for its role in cytoskeletal regulation and vascular patterning [74]. Dysregulation of SLIT1, its receptor PLXNC1, the CDC42 inhibitor SRGAP, calcium modulator PLCγ, and actin phosphatase SSH3 [75, 76] suggests impaired cytoskeletal coordination. Since many of these receptors converge on RHO family GTPases, their altered expression likely compromises RHOA-mediated actin filament remodeling and vessel stability. Collectively, these alterations point to widespread signalling imbalances that may underlie defective vascular remodeling and fragile collateral formation in MMD.

Complementing these observations, gene ontology enrichment analysis highlighted dysregulation of pathways linked to angiogenic support. Upregulation of ATP-binding cassette transporters (ABCB8, ABCA10, ABCA2) suggests metabolic adaptation to sustain neovascularization. However, concurrent disturbances in oxygen transport (HBB, HBA1), pro-angiogenic CYP4Z1, ECM regulators (CLASP1), and microtubule-associated proteins (NCKAP5) indicate a reactive angiogenic response lacking structural integrity. Of note, CYP4Z1, a cytochrome P450 monooxygenase, has been shown to enhance VEGFA expression and promote endothelial proliferation, migration, and tube formation *in vitro* and *in vivo* [77–81]. Its dysregulation in MMD therefore highlights maladaptive vascular responses to chronic ischemia.

Altogether, these findings suggest that in MMD, chronic ischemia triggers vascular remodelling and compensatory but disorganized angiogenic response, characterized by impaired cytoskeletal coordination, defective focal adhesion and PI3K-Akt signalling, and maladaptive ECM remodeling. These disrupted pathways collectively compromise vascular remodeling and contribute to the formation of structurally fragile collaterals.

### Limitations

The first major limitation of the study is the small sample size. However, given the rarity of Moyamoya disease and the limited number of patients diagnosed annually at PGIMER, this remains an inherent constraint associated with the diseases. Secondly, it was difficult to include age- and gender-matched controls, which may affect the strength of comparative analyses. This limitation is particularly relevant as most MMD patients in our cohort were from the paediatric population, making it ethically and practically challenging to obtain samples from healthy paediatric controls. Additionally, due to the limited number of cases, we were unable to stratify patients based on clinical subtypes (ischemic vs. hemorrhagic), which could have offered deeper insight into phenotype-specific mechanisms. Finally, an important limitation was the inability to perform proteomic profiling of affected vascular tissue. This arises due to the nature of the STA-MCA bypass surgery, which deliberately avoids the occluded segments to reduce surgical risk, limiting our analysis to peripheral samples that may not fully capture localized disease mechanisms.

### Future Perspective

Although several mechanisms have been implicated in Moyamoya disease, the early triggers of endothelial activation, VSMC dedifferentiation, and maladaptive vascular remodeling remain unclear. Our exosomal proteomic findings highlight disruptions in RHOA–ROCK, PI3K-Akt, Ca²⁺ signalling, and cytoskeletal regulation, suggesting that circulating vesicles may actively contribute to these early pathological events. However, it is still unknown whether these alterations originate from genetic susceptibility, chronic inflammation, altered hemodynamic, infectious stimuli, or their convergence. Future studies integrating spatial transcriptomics, single-cell profiling, multi-omics of exosomes, and advanced in vivo models will be essential to validate these pathways, pinpoint initiating drivers, and identify therapeutic targets that can modulate EC–VSMC crosstalk in early MMD.

## Conclusion

Our proteomic profiling of serum-derived exosomes demonstrates that exosomal signatures in Moyamoya disease mirror the key molecular disturbances driving its vascular pathology. We identified 213 significantly dysregulated proteins, revealing coordinated impairment in RHOA-ROCK, PI3K-Akt, focal adhesion, and Ca²⁺-dependent signalling pathways. These changes, including downregulation of RHOA, ARHGEF37, STIM1, and PLCG1, and upregulation of PRKG2, MYC, integrins, and ECM-remodelling proteins, indicate disrupted VSMC phenotypic switching, defective cytoskeletal dynamics, and impaired endothelial migration and angiogenesis. Collectively, these alterations highlight dysfunctional VSMC-EC communication and provide mechanistic insight into intimal thickening, progressive stenosis, and the formation of fragile collateral vessels in MMD. Thus, our findings support the premise that serum-derived exosomes serve as reflective biomarkers of pathogenic vascular signalling in Moyamoya disease and may offer novel targets for future therapeutic and diagnostic strategies.

## Ethics approval and consent to participate

This study was approved by the Institute Ethics Committee of PGIMER, Chandigarh, India (Ethics approval no. IEC-INT/2023/PhD-858). All the subjects were enrolled only after obtaining the written informed consent from relatives.

## Consent for publication

NA

## Availability of data and material

Data available on request.

## Author’s contribution

Conceptualization: TG, MK, AA; Sample and data collection: RB, AA; Research facilities: TG; Data Analysis: TG, RB, VD, MK; Data Interpretation: TG, RB, VD, MK, AA,; Project administration: TG; Supervision: TG; Visualization: MK, VD; Writing original draft preparation: RB, VD, MK; Writing-review and editing: RB, VD, MK, TG. All authors read and approved the final manuscript.

## Acknowledgements

The authors are highly grateful to the Council of Scientific and Industrial Research, New Delhi, India for financial assistance to R.B. in the form of Junior Research Fellowship/ Senior Research Fellowship (Award No. 09/0141(13896)/2022-EMR-1). The authors are also thankful to the subjects and their family members for the participation in this study.

## Abbreviations

MMD: Moyamoya disease
ICA: Internal carotid arteries
TIAs: Transient ischemic attacks
VSMCs: Vascular smooth muscle cells
ECM: Extracellular Matrix
MRA: Magnetic Resonance Angiography
DSA: Digital Subtraction Angiography
SDEs: Serum derived exosomes
TSG101: Tumor susceptibility gene 101
ALIX: ALG-2-interacting protein X
LAMP2: Lysosomal associated membrane protein 2
CD63: CD63 antigen
ALB: Albumin
RIPA: Radio Immunoprecipitation assay
DTT: Dithiothreitol
IAA: Iodoacetamide
PRKG2: Protein kinase cGMP-dependent 2
MYC: Myc proto-oncogene protein
RHOA: Ras homolog family member A

