## SUPPLEMENTARY for "Exosome-Derived Proteomic Signatures Highlight Pathogenic Mechanisms in Moyamoya Disease": SUPPLEMENTARY 18 APRIL 2026 - Copy.docx

| **Sr. No**  **Supplementary table 1: List of differentially expressed proteins in DIA data** | **Group** | **Accession No.** | **Protein Name** | | **Gene Symbol** | **Protein Description** | **Log2FC** | **P-value** |
| --- | --- | --- | --- | --- | --- | --- | --- | --- |
| 1. | C vs MMD | Q9GZM6 | OR8D2 | OR8D2 | | Olfactory Receptor Family 8 Subfamily D Member 2 | 10.084 | 0.01 |
| 2. | C vs MMD | Q2KHM9-1 | KIAA0753 | KIAA0753 | | Protein moonraker | 9.7742 | 0.013 |
| 3. | C vs MMD | Q7Z2Z1 | TICRR | TICRR | | TOPBP1 interacting checkpoint and replication regulator | 9.7277 | 0.019 |
| 4. | C vs MMD | P12111 | COL6A3 | COL6A3 | | Collagen type VI alpha 3 chain | 9.6916 | 0.045 |
| 5. | C vs MMD | I3L4T5 | NAA60 | NAA60 | | N-alpha-acetyltransferase 60, NatF catalytic subunit | 9.6196 | 0.027 |
| 6. | C vs MMD | Q9NVE5 | USP40 | USP40 | | Ubiquitin specific peptidase 40 | 9.5605 | 0.007 |
| 7. | C vs MMD | Q86W10 | CYP4Z1 | CYP4Z1 | | Cytochrome P450 family 4 subfamily Z member 1 | 9.4129 | 0.04 |
| 8. | C vs MMD | O14513-1 | NCKAP5 | NCKAP5 | | NCK associated protein 5 | 9.3673 | 0.001 |
| 9. | C vs MMD | Q5JVZ5 | ELMO2 | ELMO2 | | Engulfment and cell motility 2 | 9.3613 | 0.01 |
| 10. | C vs MMD | Q04637 | EIF4G1 | EIF4G1 | | Eukaryotic translation initiation factor 4 gamma 1 | 9.2913 | 0.038 |
| 11. | C vs MMD | P20711-1 | DDC | DDC | | Dopa decarboxylase | 9.2233 | 0.016 |
| 12. | C vs MMD | Q99808 | SLC29A1 | SLC29A1 | | Solute carrier family 29 member 1 | 8.8702 | 0.027 |
| 13. | C vs MMD | Q86SR1 | GLANT10 | GLANT10 | | Polypeptide N-acetylgalactosaminyltransferase 10 | 8.8204 | 0.003 |
| 14. | C vs MMD | Q9NUT2-4 | ABCB8 | ABCB8 | | ATP binding cassette subfamily B member 8 | 8.78 | 0.008 |
| 15. | C vs MMD | Q86Z20-1 | CCDC125 | CCDC125 | | Coiled-coil domain containing 125 | 8.7769 | 0.017 |
| 16. | C vs MMD | Q8NA03 | FSIP1 | FSIP1 | | Fibrous sheath interacting protein 1 | 8.7497 | 0.042 |
| 17. | C vs MMD | Q8WWZ4 | ABCA10 | ABCA10 | | ATP binding cassette subfamily A member 10 | 8.6583 | 0.008 |
| 18. | C vs MMD | K7EQL6 | USP32 | USP32 | | Ubiquitin specific peptidase 32 | 8.6276 | 0.001 |
| 19. | C vs MMD | Q96E52-1 | OMA1 | OMA1 | | OMA1 zinc metallopeptidase | 8.2402 | 0.007 |
| 20. | C vs MMD | Q13237 | PRKG2 | PRKG2 | | Protein kinase cGMP-dependent 2 | 8.2318 | 0.01 |
| 21. | C vs MMD | O60333-3 | KIF1B | KIF1B | | Kinesin family member 1B | 8.2263 | 0 |
| 22. | C vs MMD | Q13907-2 | IDI1 | IDI1 | | Isopentenyl-diphosphate delta isomerase 1 | 8.2019 | 0.041 |
| 23. | C vs MMD | Q9Y6X9-1 | MORC2 | MORC2 | | MORC family CW-type zinc finger 2 | 8.0024 | 0.016 |
| 24. | C vs MMD | K7ESA4 | ATF7 | ATF7 | | Activating transcription factor 7 | 7.838 | 0.043 |
| 25. | C vs MMD | Q92539 | LPIN2 | LPIN2 | | Lipin 2 | 7.7023 | 0.047 |
| 26. | C vs MMD | O60296 | TRAK2 | TRAK2 | | Trafficking kinesin protein 2 | 7.6571 | 0.048 |
| 27. | C vs MMD | Q3ZCT1 | ZNF260 | ZNF260 | | Zinc finger protein 260 | 7.6256 | 0.03 |
| 28. | C vs MMD | O60811 | PRAMEF2 | PRAMEF2 | | PRAME family member 2 | 7.6199 | 0.009 |
| 29. | C vs MMD | Q6SJ93-1 | FAM111B | FAM111B | | FAM111 trypsin like peptidase B | 7.6028 | 0.033 |
| 30. | C vs MMD | Q6UQ28 | PLET1 | PLET1 | | Placenta expressed transcript 1 | 7.5969 | 0.048 |
| 31. | C vs MMD | Q96N23 | CFAP54 | CFAP54 | | Cilia and flagella associated protein 54 | 7.5472 | 0.001 |
| 32. | C vs MMD | C9JPF8 | MAP6D1 | MAP6D1 | | MAP6 domain containing 1 | 7.5417 | 0.017 |
| 33. | C vs MMD | P49759 | CLK1 | CLK1 | | CDC like kinase 1 | 7.399 | 0.009 |
| 34. | C vs MMD | Q8NFD2 | ANKK1 | ANKK1 | | Ankyrin repeat and kinase domain containing 1 | 7.3785 | 0.012 |
| 35. | C vs MMD | P01106 | MYC | MYC | | MYC proto-oncogene | 7.374 | 0.002 |
| 36. | C vs MMD | A0A087X0U9 | KIAA0319 | KIAA0319 | | Dyslexia-associated protein KIAA0319 | 7.2993 | 0.037 |
| 37. | C vs MMD | Q6ZUS6-3 | CCDC149 | CCDC149 | | Coiled-coil domain containing 149 | 7.2152 | 0.009 |
| 38. | C vs MMD | P41091 | EIF2S3 | EIF2S3 | | Eukaryotic translation initiation factor 2 subunit gamma | 7.2026 | 0.022 |
| 39. | C vs MMD | Q92901 | RPL3L | RPL3L | | Ribosomal protein uL3-like | 7.2014 | 0.027 |
| 40. | C vs MMD | Q8NCP5 | ZBTB44 | ZBTB44 | | Eukaryotic translation initiation factor 2 subunit gamma | 7.0571 | 0.002 |
| 41. | C vs MMD | P49767 | VEGFC | VEGFC | | Vascular endothelial growth factor C | 7.0536 | 0.029 |
| 42. | C vs MMD | O60486 | PLXNC1 | PLXNC1 | | Plexin C1 | 7.0294 | 0.022 |
| 43. | C vs MMD | Q9Y3Y2 | CHTOP | CHTOP | | Chromatin target of PRMT1 | 6.9868 | 0.008 |
| 44. | C vs MMD | P98095 | FBLN2 | FBLN2 | | Fibulin 2 | 6.9426 | 0.031 |
| 45. | C vs MMD | Q9Y5B6 | PAXBP1 | PAXBP1 | | PAX3 and PAX7 binding protein 1 | 6.8654 | 0 |
| 46. | C vs MMD | E5RG88 | CCN4 | CCN4 | | Cellular communication network factor 4 | 6.8264 | 0.02 |
| 47. | C vs MMD | Q9UBS4 | DNAJB11 | DNAJB11 | | DnaJ heat shock protein family (Hsp40) member B11 | 6.86 | 0.047 |
| 48. | C vs MMD | Q573B4 | LCK1 | LCK1 | | LCK proto-oncogene, Src family tyrosine kinase | 6.8318 | 0.038 |
| 49. | C vs MMD | Q7Z460-1 | CLASP1 | CLASP1 | | Cytoplasmic linker associated protein 1 | 6.7821 | 0.001 |
| 50. | C vs MMD | Q96F24-1 | NRBF2 | NRBF2 | | Nuclear receptor binding factor 2 | 6.7403 | 0.028 |
| 51. | C vs MMD | Q9UJT2 | TSKS | TSKS | | Testis specific serine kinase substrate | 6.6492 | 0.037 |
| 52. | C vs MMD | Q3KRB8 | ARHGAP11B | ARHGAP11B | | Rho GTPase activating protein 11B | 6.6339 | 0.012 |
| 53. | C vs MMD | P49768 | PSEN1 | PSEN1 | | Presenilin 1 | 6.6135 | 0.017 |
| 54. | C vs MMD | A8K7K2 | NKTR | NKTR | | Natural killer cell triggering receptor | 6.6127 | 0.001 |
| 55. | C vs MMD | Q8WUY9 | DEPDC1 | DEPDC1 | | DEP domain containing 1B | 6.6044 | 0.044 |
| 56. | C vs MMD | Q86WN1-1 | FCHSD1 | FCHSD1 | | FCH and double SH3 domains 1 | 6.5783 | 0.03 |
| 57. | C vs MMD | Q9HCN3 | TMEM8A | TMEM8A | | Transmembrane protein 178A | 6.5274 | 0.029 |
| 58. | C vs MMD | Q6VMQ6 | ATF7IP | ATF7IP | | Activating transcription factor 7 interacting protein | 6.4907 | 0.044 |
| 59. | C vs MMD | E9PM06 | CHKA | CHKA | | Choline kinase alpha | 6.4883 | 0.028 |
| 60. | C vs MMD | Q13683 | ITGA7 | ITGA7 | | Integrin subunit alpha 7 | 6.3943 | 0.005 |
| 61. | C vs MMD | Q7L3S4 | ZNF771 | ZNF771 | | Zinc finger protein 771 | 6.3674 | 0.005 |
| 62. | C vs MMD | Q9H2G9-1 | BLZF1 | BLZF1 | | Basic leucine zipper nuclear factor 1 | 6.1962 | 0.012 |
| 63. | C vs MMD | Q9Y6R1-1 | SLC4A4 | SLC4A4 | | Solute carrier family 4 member 4 | 6.1933 | 0.033 |
| 64. | C vs MMD | Q8IY26 | PPAPDC2 | PPAPDC2 | | Phospholipid phosphatase 6 | 6.0649 | 0.028 |
| 65. | C vs MMD | Q5VZ89 | DENND4C | DENND4C | | DENN domain containing 4C | 6.0163 | 0.047 |
| 66. | C vs MMD | Q9Y2K3 | MYH15 | MYH15 | | Myosin heavy chain 15 | 6.0134 | 0.03 |
| 67. | C vs MMD | Q9BUP0 | EFHD1 | EFHD1 | | EF-hand domain family member D1 | 5.9688 | 0.034 |
| 68. | C vs MMD | P49792 | RANBP2 | RANBP2 | | E3 SUMO-protein ligase RanBP2 | 5.9169 | 0.038 |
| 69. | C vs MMD | Q9NZ71 | RTEL1 | RTEL1 | | Regulator of telomere elongation helicase 1 | 5.8821 | 0.034 |
| 70. | C vs MMD | P68871 | HBB | HBB | | Hemoglobin subunit beta | 5.8817 | 0.028 |
| 71. | C vs MMD | O76021 | RSL1D1 | RSL1D1 | | Ribosomal L1 domain containing 1 | 5.8359 | 0.046 |
| 72. | C vs MMD | P51451 | BLK | BLK | | BLK proto-oncogene, Src family tyrosine kinase | 5.8049 | 0.01 |
| 73. | C vs MMD | Q8NHS4-1 | CLHC1 | CLHC1 | | Clathrin heavy chain linker domain containing 1 | 5.7937 | 0.015 |
| 74. | C vs MMD | O43896 | KIF1C | KIF1C | | Kinesin family member 1B | 5.75 | 0.001 |
| 75. | C vs MMD | Q9HC36 | RNMTL1 | MRM3 | | rRNA methyltransferase 3, mitochondrial | 5.737 | 0.03 |
| 76. | C vs MMD | Q6ZP82 | CCDC141 | CCDC141 | | Coiled-coil domain containing 141 | 5.7087 | 0.019 |
| 77. | C vs MMD | A0A087WWG1 | SSH3 | SSH3 | | Slingshot protein phosphatase 3 | 5.67 | 0.009 |
| 78. | C vs MMD | Q8TF42 | UBASH3B | UBASH3B | | Ubiquitin associated and SH3 domain containing B | 5.647 | 0.043 |
| 79. | C vs MMD | Q8IZU9-2 | KIRREL3 | KIRREL3 | | Kirre like nephrin family adhesion molecule 3 | 5.6454 | 0.036 |
| 80. | C vs MMD | Q86W25 | NLRP13 | NLRP13 | | NLR family pyrin domain containing 13 | 5.5298 | 0.008 |
| 81. | C vs MMD | Q96I34 | PPP1R16A | PPP1R16A | | Protein phosphatase 2 regulatory subunit B'delta | 5.5253 | 0.044 |
| 82. | C vs MMD | Q9NQW1-1 | SEC31B | SEC31B | | SEC31 homolog B, COPII coat complex component | 5.4796 | 0.043 |
| 83. | C vs MMD | O75891 | ALDH1L1 | ALDH1L1 | | Aldehyde dehydrogenase 1 family member L1 | 5.4762 | 0.015 |
| 84. | C vs MMD | Q8TF72 | SHROOM3 | SHROOM3 | | Shroom family member 3 | 5.4535 | 0.009 |
| 85. | C vs MMD | Q9NPA3 | MID1IP1 | MID1IP1 | | MID1 interacting protein 1 | 5.3721 | 0.011 |
| 86. | C vs MMD | A0A1B0GU87 | EDDM13 | EDDM13 | | Epididymal protein 13 | 5.3543 | 0.005 |
| 87. | C vs MMD | Q96L73-1 | NSD1 | NSD1 | | Nuclear receptor binding SET domain protein 1 | 5.2571 | 0.003 |
| 88. | C vs MMD | P69905 | HBA1, HBA2 | HBA1, HBA2 | | Hemoglobin subunit alpha 2 | 5.2467 | 0.035 |
| 89. | C vs MMD | Q96QC0 | PPP1R10 | PPP1R10 | | Serine/threonine-protein phosphatase 1 regulatory subunit 10 | 5.1928 | 0.027 |
| 90. | C vs MMD | O14776-1 | TCERG1 | TCERG1 | | Transcription elongation regulator 1 | 5.0981 | 0.027 |
| 91. | C vs MMD | Q6UXM1-1 | LRIG3 | LRIG3 | | Leucine rich repeats and immunoglobulin like domains 3 | 5.0863 | 0.007 |
| 92. | C vs MMD | Q13470 | TNK1 | TNK1 | | Tyrosine kinase non receptor 1 | 5.0177 | 0.046 |
| 93. | C vs MMD | P48067 | SLC6A9 | SLC6A9 | | Solute carrier family 6 member 9 | 5.0039 | 0.031 |
| 94. | C vs MMD | Q05084 | ICA1 | ICA1 | | Islet cell autoantigen 1 | 4.9951 | 0.023 |
| 95. | C vs MMD | Q5SQ64 | LY6G6F | LY6G6F | | Lymphocyte antigen 6 family member G6F | 4.9382 | 0.015 |
| 96. | C vs MMD | Q8NAX2 | KDF1 | KDF1 | | Keratinocyte differentiation factor 1 | 4.6796 | 0.032 |
| 97. | C vs MMD | Q969P6 | TOP1MT | TOP1MT | | DNA topoisomerase I mitochondrial | 4.5794 | 0.002 |
| 98. | C vs MMD | Q9UNH6 | SNX7 | SNX7 | | Sorting nexin 7 | 4.4305 | 0.012 |
| 99. | C vs MMD | Q8NBL3 | TMEM178A | TMEM178A | | Transmembrane protein 178A | 4.3697 | 0.022 |
| 100. | C vs MMD | Q9UHQ4 | BCAP29 | BCAP29 | | B cell receptor associated protein 29 | 4.3319 | 0.038 |
| 101. | C vs MMD | Q9Y2X0-1 | MED16 | MED16 | | Mediator complex subunit 16 | 4.3012 | 0.046 |
| 102. | C vs MMD | Q9NZ08-1 | ERAP1 | ERAP1 | | Endoplasmic reticulum aminopeptidase 1 | 4.2728 | 0.015 |
| 103. | C vs MMD | Q68CQ1-7 | MROH7 | MROH7 | | Maestro heat like repeat family member 7 | 4.2689 | 0.02 |
| 104. | C vs MMD | Q5TG30 | ARHGAP40 | ARHGAP40 | | Rho GTPase-activating protein 40 | 4.2481 | 0.049 |
| 105. | C vs MMD | A0A087WXC8 | CYCL1 | CYCL1 | | Cylicin 1 | 4.0328 | 0.017 |
| 106. | C vs MMD | Q96IW2 | SHD | SHD | | Src homology 2 domain containing transforming protein D | 4.0116 | 0.02 |
| 107. | C vs MMD | E5RGQ2 | AHRR | AHRR | | Aryl hydrocarbon receptor repressor | 3.9419 | 0.024 |
| 108. | C vs MMD | Q8NDB2 | BANK1 | BANK1 | | B cell scaffold protein with ankyrin repeats 1 | 3.7092 | 0.004 |
| 109. | C vs MMD | A0A075B7B7 | Uncharacterized Protein | Uncharacterized Protein | | Uncharacterized Protein | 3.6007 | 0.047 |
| 110. | C vs MMD | Q7RTT3 | SSX9P | SSX9P | | Putative protein SSX9 | 3.3868 | 0.022 |
| 111. | C vs MMD | O96028-1 | NSD2 | NSD2 | | Nuclear receptor binding SET domain protein 2 | 3.3215 | 0.036 |
| 112. | C vs MMD | F5H2G6 | ACSF3 | ACSF3 | | Acyl-CoA synthetase family member 3 | 2.8806 | 0.047 |
| 113. | C vs MMD | P01871 | IGHM | IGHM | | Immunoglobulin heavy constant mu | 2.7342 | 0.003 |
| 114. | C vs MMD | Q9BZC7 | ABCA2 | ABCA2 | | ATP binding cassette subfamily A member 2 | 2.6135 | 0.034 |
| 115. | C vs MMD | P02787 | TF | TF | | Transferrin | 2.3554 | 0.047 |
| 116. | C vs MMD | A8MX80 | Uncharacterized Protein | Uncharacterized Protein | | Uncharacterized Protein | 1.769 | 0.048 |
| 117. | C vs MMD | Q2M385 | MPEG1 | MPEG1 | | Macrophage expressed 1 | 1.6769 | 0.017 |
| 118. | C vs MMD | Q8N103-1 | TAGAP | TAGAP | | T cell activation RhoGTPase activating protein | 0.95315 | 0.041 |
| 119. | C vs MMD | O60307 | MAST3 | MAST3 | | Microtubule associated serine/threonine kinase 3 | -1.055 | 0.04 |
| 120. | C vs MMD | Q16531 | DDB1 | DDB1 | | Damage specific DNA binding protein 1 | -1.2896 | 0.028 |
| 121. | C vs MMD | Q2LD37-1 | BLTP1 | BLTP1 | | Bridge-like lipid transfer protein family member 1 | -1.4081 | 0.001 |
| 122. | C vs MMD | Q8NI17-10 | IL31RA | IL31RA | | Interleukin 31 receptor A | -1.5351 | 0.043 |
| 123. | C vs MMD | Q86UW7 | CADPS2 | CADPS2 | | Calcium dependent secretion activator 2 | -1.7248 | 0.038 |
| 124. | C vs MMD | Q13586 | STIM1 | STIM1 | | Stromal interaction molecule 1 | -1.8666 | 0.004 |
| 125. | C vs MMD | P02458-2 | COL2A1 | COL2A1 | | Collagen type II alpha 1 chain | -2.231 | 0.021 |
| 126. | C vs MMD | Q5JRM2 | CXorf66 | CXorf66 | | Chromosome X open reading frame 66 | -2.4455 | 0.003 |
| 127. | C vs MMD | H7C4W0 | ASIC3 | ASIC3 | | Acid sensing ion channel subunit 3 | -2.6068 | 0.038 |
| 128. | C vs MMD | Q14003 | KCNC3 | KCNC3 | | Potassium voltage-gated channel subfamily C member 3 | -2.7338 | 0.038 |
| 129. | C vs MMD | P20929 | NEB | NEB | | Nebulin | -2.8122 | 0.016 |
| 130. | C vs MMD | Q6NSJ5 | LRRC8E | LRRC8E | | Leucine rich repeat containing 8 VRAC subunit E | -3.0391 | 0.043 |
| 131. | C vs MMD | Q9P035 | HACD3 | HACD3 | | 3-hydroxyacyl-CoA dehydratase 3 | -3.1266 | 0.001 |
| 132. | C vs MMD | Q8NCW0 | KREMEN2 | KREMEN2 | | Kringle containing transmembrane protein 2 | -3.2596 | 0.031 |
| 133. | C vs MMD | P30825 | SLC7A1 | SLC7A1 | | High affinity cationic amino acid transporter 1 | -3.2662 | 0.009 |
| 134. | C vs MMD | Q9H4Q3 | PRDM13 | PRDM13 | | PR/SET domain 13 | -4.0123 | 0.033 |
| 135. | C vs MMD | P0C7U3 | ZDHHC11B | ZDHHC11B | | Predicted to enable protein-cysteine S-palmitoyltransferase activity | -4.114 | 0.021 |
| 136. | C vs MMD | E5RFU3 | DCTN6 | DCTN6 | | Dynactin subunit 6 | -4.244 | 0.025 |
| 137. | C vs MMD | Q8IY50 | SLC35F3 | SLC35F3 | | Solute carrier family 35 member F3 | -4.2751 | 0.047 |
| 138. | C vs MMD | H3BR70 | Uncharacterized Protein | Uncharacterized Protein | | Uncharacterized Protein | -4.3098 | 0.032 |
| 139. | C vs MMD | O75044 | SRGAP2 | SRGAP2 | | SLIT-ROBO Rho GTPase activating protein 2 | -4.3511 | 0.038 |
| 140. | C vs MMD | Q7Z434-3 | MAVS | MAVS | | Mitochondrial antiviral signaling protein | -4.3712 | 0.038 |
| 141. | C vs MMD | Q969V3 | NCLN | NCLN | | Nicalin | -4.3811 | 0.012 |
| 142. | C vs MMD | Q7Z4T9-7 | CFAP91 | CFAP91 | | Cilia and flagella associated protein 91 | -4.3999 | 0.022 |
| 143. | C vs MMD | P09038-1 | FGF2 | FGF2 | | Fibroblast growth factor 2 | -4.4455 | 0.032 |
| 144. | C vs MMD | Q96MC6 | HIAT1 | MFSD14A | | Hippocampus abundant transcript 1 protein | -4.4977 | 0.017 |
| 145. | C vs MMD | Q96IG2-1 | FBXL20 | FBXL20 | | F-box and leucine rich repeat protein 20 | -4.5088 | 0.001 |
| 146. | C vs MMD | Q96KG9 | SCYL1 | SCYL1 | | SCY1 like pseudokinase 1 | -4.5511 | 0.043 |
| 147. | C vs MMD | Q9Y6Z7 | COLEC10 | COLEC10 | | Collectin subfamily member 10 | -4.6166 | 0.027 |
| 148. | C vs MMD | Q8NHW4-10 | CCL4L1, CCL4L2 | CCL4L1, CCL4L2 | | C-C motif chemokine ligand 4 like 2 | -4.6431 | 0.001 |
| 149. | C vs MMD | Q93073-1 | SECISBP2L | SECISBP2L | | SECIS binding protein 2 like | -4.6473 | 0.034 |
| 150. | C vs MMD | P15088 | CPA3 | CPA3 | | Carboxypeptidase A3 | -4.6715 | 0.038 |
| 151. | C vs MMD | P61586 | RHOA | RHOA | | Ras homolog family member A | -4.6779 | 0.033 |
| 152. | C vs MMD | P82933 | MRPS9 | MRPS9 | | Mitochondrial ribosomal protein L16 | -4.7355 | 0.03 |
| 153. | C vs MMD | A8MYP8 | ODF3B | ODF3B | | Ciliary microtubule associated protein 1B | -4.8017 | 0.037 |
| 154. | C vs MMD | H7C5Q9 | PPP2R5D | PPP2R5D | | Serine/threonine-protein phosphatase 2A 56 kDa regulatory subunit delta isoform | -4.8254 | 0.041 |
| 155. | C vs MMD | Q9C098 | DCLK3 | DCLK3 | | Doublecortin like kinase 3 | -4.828 | 0.043 |
| 156. | C vs MMD | Q9BQ39 | DDX50 | DDX50 | | DExD-box helicase 50 | -4.8821 | 0.001 |
| 157. | C vs MMD | Q8IV35 | WDR49 | WDR49 | | Cillia & flagella associated portein | -4.9828 | 0.029 |
| 158. | C vs MMD | Q8IUD2-1 | ERC1 | ERC1 | | ELKS/RAB6-interacting/CAST family member 1 | -4.9968 | 0.043 |
| 159. | C vs MMD | Q13905-1 | RAPGEF1 | RAPGEF1 | | Rap guanine nucleotide exchange factor 1 | -5.0084 | 0.009 |
| 160. | C vs MMD | Q7Z7A3 | CTU1 | CTU1 | | Cytosolic thiouridylase subunit 1 | -5.0929 | 0.021 |
| 161. | C vs MMD | Q9H7D7 | WDR26 | WDR26 | | WD repeat domain 26 | -5.1294 | 0.008 |
| 162. | C vs MMD | Q6ZV29-4 | PNPLA7 | PNPLA7 | | Patatin-like phospholipase domain-containing protein 7 | -5.1376 | 0.038 |
| 163. | C vs MMD | Q9Y2D4 | EXOC6B | EXOC6B | | Exocyst complex component 6B | -5.167 | 0.034 |
| 164. | C vs MMD | G3V3G0 | PROX2 | PROX2 | | Prospero homeobox 2 | -5.1964 | 0.01 |
| 165. | C vs MMD | B2RTY4-4 | MYO9A | MYO9A | | Myosin IXA | -5.2108 | 0.003 |
| 166. | C vs MMD | Q9BXI9 | C1QTNF6 | C1QTNF6 | | C1q and TNF related 6 | -5.211 | 0.015 |
| 167. | C vs MMD | Q9H8W4 | PLEKHF2 | PLEKHF2 | | Pleckstrin homology domain-containing family F member 2 | -5.2566 | 0.023 |
| 168. | C vs MMD | P19174-1 | PLCG1 | PLCG1 | | Phospholipase C gamma 1 | -5.2861 | 0.013 |
| 169. | C vs MMD | P0CZ25 | DNAH10OS | DNAH10OS | | Uncharacterized protein DNAH10OS | -5.3074 | 0.041 |
| 170. | C vs MMD | Q8NI35-5 | PATJ | PATJ | | InaD-like protein | -5.3269 | 0.029 |
| 171. | C vs MMD | Q9BYD6 | MRLP1 | MRLP1 | | C-type lectin domain family 18 member B (CLEC18B) | -5.3628 | 0.027 |
| 172. | C vs MMD | Q6V1P9 | DCHS2 | DCHS2 | | Dachsous cadherin-related 2 | -5.366 | 0.044 |
| 173. | C vs MMD | P62316 | SNRPD2 | SNRPD2 | | Small nuclear ribonucleoprotein D2 polypeptide | -5.4456 | 0.009 |
| 174. | C vs MMD | Q8WVJ2 | NUDCD2 | NUDCD2 | | NudC domain containing 2 | -5.4689 | 0.036 |
| 175. | C vs MMD | Q03924 | ZNF117 | ZNF117 | | Zinc finger protein 117 | -5.4787 | 0.049 |
| 176. | C vs MMD | A0A1B0GTP3 | CCDC196 | CCDC196 | | Coiled coil Domain containing 196 | -5.4934 | 0.034 |
| 177. | C vs MMD | Q6ZMN7-1 | PDZRN4 | PDZRN4 | | PDZ domain-containing RING finger protein 4 | -5.5399 | 0.024 |
| 178. | C vs MMD | C9JUZ5 | PEX5L | PEX5L | | PEX5-related protein | -5.5505 | 0.033 |
| 179. | C vs MMD | P48549-2 | KCNJ3 | KCNJ3 | | Potassium inwardly rectifying channel subfamily J member 3 | -5.6059 | 0.019 |
| 180. | C vs MMD | Q6Q4G3 | LVRN | LVRN | | Laeverin | -5.7603 | 0.033 |
| 181. | C vs MMD | Q9H7T0 | CATSPERB | CATSPERB | | Cation channel sperm associated auxiliary subunit beta | -5.7788 | 0.001 |
| 182. | C vs MMD | A1IGU5 | ARHGEF37 | ARHGEF37 | | Rho guanine nucleotide exchange factor 37 | -5.8865 | 0.009 |
| 183. | C vs MMD | P61925 | PKIA | PKIA | | cAMP-dependent protein kinase inhibitor alpha | -5.9005 | 0.033 |
| 184. | C vs MMD | Q5H9U9-1 | DDX60L | DDX60L | | DExD/H-box 60 like | -5.9432 | 0.031 |
| 185. | C vs MMD | P56181-2 | NUDFV3 | NUDFV3 | | NADH ubiquinone oxidoreductase subunit V3 | -6.0414 | 0.038 |
| 186. | C vs MMD | Q9NXZ1 | SAGE1 | SAGE1 | | Sarcoma antigen 1 | -6.072 | 0.007 |
| 187. | C vs MMD | F8VXH5 | MTERF2 | MTERF2 | | Mitochondrial transcription termination factor 2 | -6.1014 | 0 |
| 188. | C vs MMD | Q17RY0-1 | CPEB4 | CPEB4 | | Cytoplasmic polyadenylation element binding protein 4 | -6.2028 | 0.009 |
| 189. | C vs MMD | Q8IUZ0-2 | LRRC49 | LRRC49 | | Leucine rich repeat containing 49 | -6.4305 | 0.049 |
| 190. | C vs MMD | E9PLF2 | SLC43A3 | SLC43A3 | | Solute carrier family 43 member 3 | -6.4368 | 0.048 |
| 191. | C vs MMD | Q8IV53-1 | DENND1C | DENND1C | | DENN domain containing 1C | -6.4498 | 0.048 |
| 192. | C vs MMD | C9J1X3 | TNK2 | TNK2 | | Tyrosine kinase non receptor 2 | -6.4793 | 0.021 |
| 193. | C vs MMD | P46782 | RPS5 | RPS5 | | Ribosomal protein S5 | -6.5053 | 0.04 |
| 194. | C vs MMD | E9PNU7 | GPR68 | GPR68 | | G protein-coupled receptor 68 | -6.7278 | 0.024 |
| 195. | C vs MMD | P12270 | TPR | TPR | | Translocated promoter region, nuclear basket protein | -6.9779 | 0.002 |
| 196. | C vs MMD | E5RIK2 | DNM3 | DNM3 | | Dynamin 3 | -6.9889 | 0.045 |
| 197. | C vs MMD | Q9NRG7-1 | Uncharacterized Protein | Uncharacterized Protein | | Uncharacterized Protein | -7.0174 | 0.049 |
| 198. | C vs MMD | Q03001-3 | DST | DST | | Dystonin | -7.1305 | 0.034 |
| 199. | C vs MMD | Q96G46 | DUS3L | DUS3L | | Dihydrouridine synthase 3 like | -7.1479 | 0.006 |
| 200. | C vs MMD | A0A2R8Y5B5 | SLC19A2 | SLC19A2 | | Solute carrier family 19 member 2 | -7.4232 | 0.047 |
| 201. | C vs MMD | U3KQL1 | PDCD2L | PDCD2L | | Programmed cell death 2 like | -7.535 | 0.032 |
| 202. | C vs MMD | P04745 | AMY1A | AMY1A | | Alpha-amylase 1A | -7.5615 | 0.045 |
| 203. | C vs MMD | Q8TAT5 | NEIL3 | NEIL3 | | Nei like DNA glycosylase 3 | -7.7866 | 0.004 |
| 204. | C vs MMD | Q9NX20 | MRPL16 | MRPL16 | | Mitochondrial ribosomal protein L1 | -7.905 | 0.014 |
| 205. | C vs MMD | Q9UBJ2 | ABCD2 | ABCD2 | | ATP binding cassette subfamily D member 2 | -8.082 | 0.033 |
| 206. | C vs MMD | P62140 | PPP1CB | PPP1CB | | Serine/threonine-protein phosphatase PP1-beta catalytic subunit | -8.0833 | 0.014 |
| 207. | C vs MMD | P28221 | HTR1D | HTR1D | | 5-hydroxytryptamine receptor 1D | -8.3432 | 0.042 |
| 208. | C vs MMD | P54756 | EPAH5 | EPAH5 | | EPH receptor A5 | -8.5743 | 0.015 |
| 209. | C vs MMD | C9J5N1 | PTGES3L-AARSD1 | AARSD1 | | Alanyl-tRNA editing protein Aarsd1 | -8.5848 | 0.034 |
| 210. | C vs MMD | Q5T6F0 | DCAF12 | DCAF12 | | DDB1 and CUL4 associated factor 12 | -9.0365 | 0.013 |
| 211. | C vs MMD | O75093-1 | SLIT1 | SLIT1 | | Slit guidance ligand 1 | -9.2521 | 0.042 |
| 212. | C vs MMD | O15530-1 | PDPK1 | PDPK1 | | 3-phosphoinositide dependent protein kinase 1 | -9.2609 | 0.045 |
| 213. | C vs MMD | Q13434 | MKRN4P | MKRN4P | | Putative E3 ubiquitin-protein ligase makorin-4 | -10.527 | 0.002 |

**Table 2: Gene Ontology (GO) including Biological Processes (BP), Molecular Function (MF) and Cellular Component (CC) enrichment analysis for the Upregulated differential expressed proteins in DIA data (Top 10 terms in C vs MMD comparison group)**

| **Sr. No** | **Group** | **GO id** | **GO Terms** | **Hits** | **Total** | **Odds Ratio** | **P-value** | **Class** |
| --- | --- | --- | --- | --- | --- | --- | --- | --- |
| 1 | C vs MMD | GO:0030705 | Cytoskeleton-Dependent Intracellular Transport | 3 | 213 | 27.5125 | 0.0003 | BP |
| 2 | C vs MMD | GO:0010941 | Regulation of Cell Death | 4 | 213 | 12.7614 | 0.0004 | BP |
| 3 | C vs MMD | GO:0019755 | One carbon compound transport | 3 | 213 | 16.3248 | 0.0011 | BP |
| 4 | C vs MMD | GO:0032204 | Regulation Of Telomere Maintenance | 3 | 213 | 14.9233 | 0.0014 | BP |
| 5 | C vs MMD | GO:0015671 | Oxygen Transport | 2 | 213 | 43.2065 | 0.0015 | BP |
| 6 | C vs MMD | GO:0008089 | Anterograde Axonal Transport | 3 | 213 | 14.508 | 0.0015 | BP |
| 7 | C vs MMD | GO:0043085 | Positive Regulation Of Catalytic Activity | 3 | 213 | 14.1152 | 0.0016 | BP |
| 8 | C vs MMD | GO:0015670 | Carbon Dioxide Transport | 2 | 213 | 38.4039 | 0.0018 | BP |
| 9 | C vs MMD | GO:0010942 | Positive Regulation Of Cell Death | 3 | 213 | 11.1064 | 0.0031 | BP |
| 10 | C vs MMD | GO:0015669 | Gas Transport | 2 | 213 | 26.5819 | 0.0034 | BP |
| 11 | C vs MMD | GO:0042799 | Histone H4K20 Methyltransferase Activity | 2 | 213 | 69.1409 | 0.0007 | MF |
| 12 | C vs MMD | GO:0004715 | Non-Membrane Spanning Protein Tyrosine Kinase | 3 | 213 | 15.8293 | 0.0012 | MF |
|  |  |  | Activity |  |  |  |  |  |
| 13 | C vs MMD | GO:0046975 | Histone H3K36 Methyltransferase Activity | 2 | 213 | 34.5617 | 0.0022 | MF |
| 14 | C vs MMD | GO:0140939 | Histone H4 Methyltransferase Activity | 2 | 213 | 31.4182 | 0.0025 | MF |
| 15 | C vs MMD | GO:0005524 | ATP Binding | 6 | 213 | 3.8828 | 0.006 | MF |
| 16 | C vs MMD | GO:0016018 | Cyclosporin A Binding | 2 | 213 | 16.4489 | 0.0079 | MF |
| 17 | C vs MMD | GO:0051019 | Mitogen-Activated Protein Kinase Binding | 2 | 213 | 16.4489 | 0.0079 | MF |
| 18 | C vs MMD | GO:0032559 | Adenyl Ribonucleotide Binding | 6 | 213 | 3.493 | 0.0097 | MF |
| 19 | C vs MMD | GO:0046966 | Nuclear Thyroid Hormone Receptor Binding | 2 | 213 | 13.8143 | 0.0108 | MF |
| 20 | C vs MMD | GO:0140938 | Histone H3 Methyltransferase Activity | 2 | 213 | 10.1529 | 0.0188 | MF |
| 21 | C vs MMD | GO:0071682 | Endocytic Vesicle Lumen | 2 | 213 | 18.1822 | 0.0066 | CC |
| 22 | C vs MMD | GO:0031594 | Neuromuscular Junction | 2 | 213 | 9.3283 | 0.0218 | CC |
| 23 | C vs MMD | GO:0005874 | Microtubule | 4 | 213 | 3.8531 | 0.0234 | CC |
| 24 | C vs MMD | GO:0043190 | ATP-binding Cassette (ABC) Transporter Complex | 1 | 213 | 42.8427 | 0.0289 | CC |
| 25 | C vs MMD | GO:0098533 | ATPase Dependent Transmembrane Transport Complex | 1 | 213 | 34.2724 | 0.0346 | CC |
| 26 | C vs MMD | GO:0072357 | PTW/PP1 Phosphatase Complex | 2 | 213 | 34.2724 | 0.0346 | CC |
|  |  |  | Extracellular Membrane-Bounded Organelle |  |  |  |  |  |
| 27 | C vs MMD | GO:006501 | Extracellular Membrane-Bounded Organelle | 2 | 213 | 6.2697 | 0.044 | CC |
| 28 | C vs MMD | GO:0005801 | cis-Golgi Network | 2 | 213 | 5.9445 | 0.0482 | CC |
| 29 | C vs MMD | GO:1903561 | Extracellular Vesicle | 2 | 213 | 5.9445 | 0.0482 | CC |

**Table 3: Gene Ontology (GO) including Biological Processes (BP), Molecular Function (MF) and Cellular Component (CC) enrichment analysis for the Downregulated differential expressed proteins in DIA data (Top 10 terms in C vs MMD comparison group)**

| **Sr. No** | **Group** | **GO id** | **GO Terms** | **Hits** | **Total** | **Odds Ratio** | **P-value** | **Class** |
| --- | --- | --- | --- | --- | --- | --- | --- | --- |
| 1 | C vs MMD | GO:1905564 | Positive Regulation of Vascular Endothelial | 3 | 213 | 73.6963 | 0 | BP |
| 2 | C vs MMD | GO:0010941 | Cell Proliferation | 4 | 213 | 12.7614 | 0.0004 | BP |
| 2 | C vs MMD | GO:0035089 | Establishment Of Apical/Basal Cell Polarity | 3 | 213 | 60.2909 | 0 | BP |
| 3 | C vs MMD | GO:1905562 | Regulation Of Vascular Endothelial Cell | 3 | 213 | 44.2044 | 0.0001 | BP |
| 5 | C vs MMD | GO:0015671 | Proliferation | 2 | 213 | 43.2065 | 0.0015 | BP |
| 4 | C vs MMD | GO:0070849 | Response To Epidermal Growth Factor | 3 | 213 | 24.5432 | 0.0004 | BP |
| 5 | C vs MMD | GO:0060193 | Positive Regulation of Lipase Activity | 2 | 213 | 62.4804 | 0.0008 | BP |
| 6 | C vs MMD | GO:0045197 | Establishment Or Maintenance of Epithelial | 3 | 213 | 17.9009 | 0.0008 | BP |
| 9 | C vs MMD | GO:0010942 | Cell Apical/Basal Polarity | 3 | 213 | 11.1064 | 0.0031 | BP |
| 7 | C vs MMD | GO:0045198 | Establishment Of Epithelial Cell Apical/Basal | 2 | 213 | 54.6676 | 0.0009 | BP |
| 11 | C vs MMD | GO:0042799 | Polarity | 2 | 213 | 69.1409 | 0.0007 | MF |
| 8 | C vs MMD | GO:0043536 | Positive Regulation of Blood Vessel Endothelial | 3 | 213 | 15.3984 | 0.0013 | BP |
|  |  |  | Cell Migration |  |  |  |  |  |
| 9 | C vs MMD | GO:1904018 | Positive Regulation of Vasculature Development | 4 | 213 | 8.902 | 0.0014 | BP |
| 10 | C vs MMD | GO:0007264 | Small GTPase Mediated Signal Transduction | 4 | 213 | 8.2393 | 0.0019 | BP |
| 11 | C vs MMD | GO:0051010 | Microtubule Plus-End Binding | 2 | 213 | 25.7143 | 0.0035 | MF |
| 12 | C vs MMD | GO:0042056 | Chemoattractant Activity | 2 | 213 | 13.2361 | 0.0115 | MF |
| 13 | C vs MMD | GO:0030695 | GTPase Regulator Activity | 6 | 213 | 3.2155 | 0.0142 | MF |
| 14 | C vs MMD | GO:0003684 | Damaged DNA Binding | 2 | 213 | 10.1528 | 0.0186 | MF |
| 15 | C vs MMD | GO:0019843 | rRNA Binding | 2 | 213 | 9.9216 | 0.0194 | MF |
| 16 | C vs MMD | GO:0005167 | Neurotrophin TRK Receptor Binding | 1 | 213 | 54.0842 | 0.023 | MF |
| 17 | C vs MMD | GO:0005168 | Neurotrophin TRKA Receptor Binding | 1 | 213 | 54.0842 | 0.023 | MF |
| 18 | C vs MMD | GO:0018812 | 3-hydroxyacyl-CoA Dehydratase Activity | 1 | 213 | 43.2652 | 0.0276 | MF |
| 19 | C vs MMD | GO:0042289 | MHC Class II Protein Binding | 1 | 213 | 43.2652 | 0.0276 | MF |
| 20 | C vs MMD | GO:0048495 | Roundabout Binding | 1 | 213 | 43.2652 | 0.0276 | MF |
| 21 | C vs MMD | GO:0031903 | Microbody Membrane | 3 | 213 | 13.238 | 0.0019 | CC |
| 22 | C vs MMD | GO:0005778 | Peroxisomal Membrane | 3 | 213 | 13.238 | 0.0019 | CC |
| 23 | C vs MMD | GO:0080008 | Cul4-RING E3 Ubiquitin Ligase Complex | 2 | 213 | 13.2361 | 0.0115 | CC |
| 24 | C vs MMD | GO:0005777 | Peroxisome | 3 | 213 | 5.2331 | 0.0223 | CC |
| 25 | C vs MMD | GO:0032541 | Cortical Endoplasmic Reticulum | 1 | 213 | 54.0842 | 0.023 | CC |
| 26 | C vs MMD | GO:0072357 | PTW/PP1 Phosphatase Complex | 1 | 213 | 43.2652 | 0.0276 | CC |

**Table 4: KEGG pathway enrichment analysis for the differential expressed proteins in DIA data (Top 20 terms).**

| **S. No** | **Groups** | **KEGG ID** | **Pathways** | **List Hits** | **List Total** | **Odds Ratio** |
| --- | --- | --- | --- | --- | --- | --- |
| 1. | C vs MMD | hsa05143 | African trypanosomiasis ↑ | 3 | 213 | 15.3630 |
| 2. | C vs MMD | hsa02010 | ABC transporters ↑ | 3 | 213 | 12.4317 |
| 3. | C vs MMD | hsa05144 | Malaria ↑ | 3 | 213 | 11.1064 |
| 4. | C vs MMD | hsa00310 | Lysine degradation ↑ | 2 | 213 | 5.6513 |
| 5. | C vs MMD | hsa04310 | Wnt signaling pathway ↑ | 3 | 213 | 3.1837 |
| 6. | C vs MMD | hsa04512 | ECM-receptor interaction ↑ | 2 | 213 | 4.0034 |
| 7. | C vs MMD | hsa00360 | Phenylalanine metabolism ↑ | 1 | 213 | 10.7042 |
| 8. | C vs MMD | hsa03013 | RNA transport ↑ | 3 | 213 | 2.8329 |
| 9. | C vs MMD | hsa01212 | Fatty acid biosynthesis ↑ | 1 | 213 | 10.0740 |
| 10. | C vs MMD | hsa00670 | One carbon pool by folate ↑ | 1 | 213 | 9.0127 |
| 11. | C vs MMD | hsa04360 | Axon guidance ↓ | 5 | 213 | 6.3335 |
| 12. | C vs MMD | hsa04722 | Neurotrophin signaling pathway ↓ | 4 | 213 | 7.7350 |
| 13. | C vs MMD | hsa04510 | Focal adhesion ↓ | 5 | 213 | 5.7140 |
| 14. | C vs MMD | hsa05205 | Proteoglycans in cancer ↓ | 5 | 213 | 5.5986 |
| 15. | C vs MMD | hsa04750 | Inflammatory mediator regulation of TRP channels ↓ | 3 | 213 | 6.9516 |
| 16. | C vs MMD | hsa04972 | Pancreatic secretion ↓ | 3 | 213 | 6.6694 |
| 17. | C vs MMD | hsa04660 | T cell receptor signaling pathway ↓ | 3 | 213 | 6.5366 |
| 18. | C vs MMD | hsa04064 | NF-kappa B signaling pathway ↓ | 3 | 213 | 6.5366 |
| 19. | C vs MMD | hsa04015 | Rap1 signaling pathway ↓ | 4 | 213 | 4.2982 |
| 20. | C vs MMD | hsa04071 | Sphingolipid signaling pathway ↓ | 3 | 213 | 5.6871 |
